# Multi-omic network analysis identified betacellulin as a novel target of omega-3 fatty acid attenuation of western diet-induced nonalcoholic steatohepatitis

**DOI:** 10.1101/2022.10.03.510635

**Authors:** Jyothi Padiadpu, Manuel Garcia-Jaramillo, Nolan K. Newman, Jacob W. Pederson, Richard Rodrigues, Zhipeng Li, Sehajvir Singh, Philip Monnier, Giorgio Trinchieri, Kevin Brown, Amiran K. Dzutsev, Natalia Shulzhenko, Donald B. Jump, Andrey Morgun

**Author notes:** Co-senior correspondence to, or. These authors contributed equally to this work (Manuel Garcia-Jaramillo, Nolan K. Newman, Jacob W. Pederson).

## Abstract

Clinical and preclinical studies have established that supplementing diets with ω3 polyunsaturated fatty acids (PUFA) can reduce hepatic dysfunction in nonalcoholic steatohepatitis (NASH). Herein, we used multi-omic network analysis to unveil novel mechanistic targets of ω3 PUFA effects in a preclinical mouse model of western diet induced NASH. After identifying critical molecular processes responsible for the effects of ω3 PUFA, we next performed meta-analysis of human liver cancer transcriptomes and uncovered betacellulin as a key EGFR-binding protein that was induced in liver cancer and downregulated by ω3 PUFAs in animals with NASH. We then confirmed that betacellulin acts by promoting proliferation of quiescent hepatic stellate cells, stimulating transforming growth factor–β2 and increasing collagen production. When used in combination with TLR2/4 agonists, betacellulin upregulated integrins in macrophages thereby potentiating inflammation and fibrosis. Taken together, our results suggest that suppression of betacellulin is one of the key mechanisms associated with anti-inflammatory and antifibrotic effects of ω3 PUFA during NASH.

**Synopsis:** 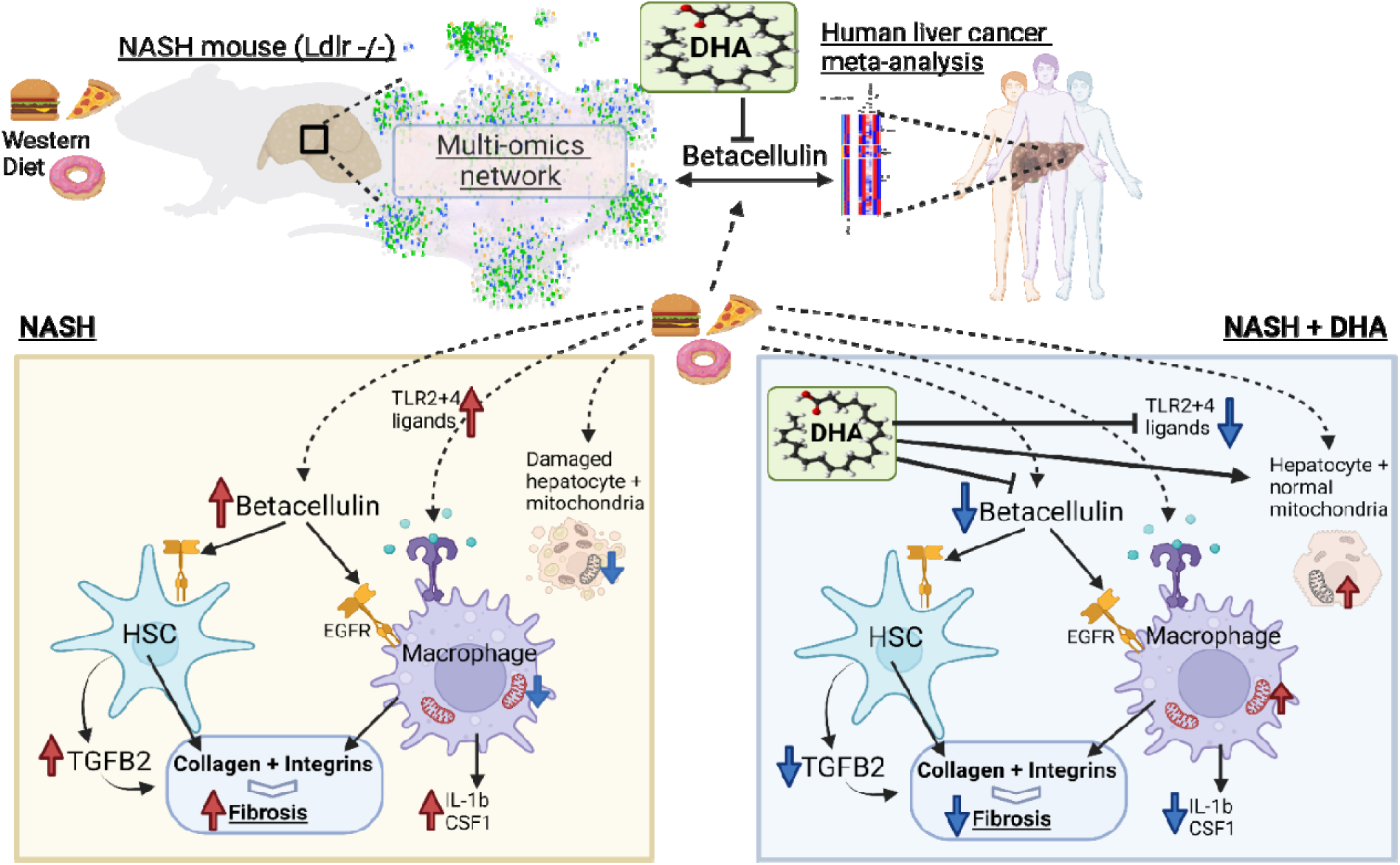

- Multi-omic network analysis points to mitochondrial cardiolipin precursors as candidate key lipids whereby ω3 fatty acids restore mitochondrial functioning.
- Multi-omic network analysis suggests betacellulin (BTC) as one of the key mediators of NASH suppressed by ω3 polyunsaturated fatty acids.
- Reduction of liver fibrosis by omega-3 fatty acids (especially by docosahexaenoic acid, DHA) is accomplished by simultaneous inhibition of betacellulin and TLR agonists.
- BTC promotes collagen production and induces TGFB2 in hepatic stellate cells.
- BTC together with TLR2/4 agonists stimulate expression of integrins in macrophages.
- DHA suppresses BTC-EGFR pathway in NASH animal model potentially preventing progression to hepatocellular carcinoma.

## Introduction

Metabolic diseases associated with obesity have increased to epidemic proportions in recent years and are one of the leading causes of morbidity and mortality (Dufour *et al*, 2022; Konerman *et al*, 2018). Metabolic-associated fatty liver disease (MAFLD) or nonalcoholic steatohepatitis (NASH), type 2 diabetes and cardiovascular diseases are all associated with obesity and a sedentary lifestyle (Lazarus *et al*, 2022; Marjot *et al*, 2020). About 80 million adults and 13 million children are obese in the US alone. Among these, 60 percent of patients with body mass index greater than 30 have evidence of liver steatosis (Dufour *et al*., 2022; Jump *et al*, 2015). NASH is a progressive form of nonalcoholic fatty liver disease (NAFLD) and is a major risk factor for cirrhosis, hepatocellular carcinoma (HCC) and liver failure. While treatments to manage the co-morbidities associated with NASH, i.e., obesity and type 2 diabetes are available, NASH has no specific FDA-approved treatment. Thus, lifestyle modifications and dietary interventions are the current options available to NASH patients. Most if not all drugs which targeted individual molecules or specific pathways have failed to significantly improve the NASH patient (Ampuero *et al*, 2022; Dufour *et al*., 2022; Neuschwander-Tetri, 2020; Pfister *et al*, 2021). This strategy that has been effective in the treatment of other diseases might not be adequate for NAFLD/NASH therapy because it does not address entirely the complexity of this disease and may miss the master regulators involved in disease onset and progression.

Omega-3 polyunsaturated fatty acids (PUFA) are known to be consistently lower in livers of NASH patients when compared to healthy patients or patients with benign steatosis (Burke *et al*, 1999; Fridén *et al*, 2021). This prompted us to hypothesize that dietary supplementation with ω3 PUFA would restore liver functions. Indeed, this strategy was very successful in a preclinical mouse model, not only reducing liver steatosis but also in attenuating western diet-induced hepatic fibrosis(Depner *et al*, 2013a; Depner *et al*, 2013b; Jump *et al*., 2015). Moreover, ω3 fatty acid treatment of children and adults with NAFLD have demonstrated these dietary lipids reduce hepatic steatosis and hepatic injury (Iannelli *et al*, 2013; Spooner & Jump, 2019). While it is well established that ω3 PUFA have the capacity to regulate hepatic mechanisms controlling fatty acid synthesis and oxidation, as well as inflammation, it is less clear if these pathways form the extent of ω3 fatty acid regulation of hepatic function (Jump *et al*, 2018). Despite several studies demonstrating the therapeutic effects of ω3 PUFA in NAFLD/NASH models the mechanism of action has been elusive. Nevertheless, it is important to note that many studies demonstrated diverse effects of ω3 in the liver ranging from immune-modulatory activities(Gutiérrez *et al*, 2019) and improvement of oxidative stress (Yang *et al*, 2019) to structural effects related to incorporation of phospholipids into the mitochondrial membrane, altogether with potentially positive impact on liver function. In addition to the effects of ω3 PUFA on hepatic cells, recent studies suggest that ω3 PUFA can alter gut microbiota, a known player in the pathogenesis of NAFLD/NASH (Watson *et al*, 2018). Furthermore, preclinical and clinical studies demonstrated that docosahexaenoic acid (DHA, 22:6, ω3) might be a more efficient agent than eicosapentaenoic acid (EPA, 20:5, ω3) in preventing and treating NAFLD (Depner *et al*., 2013a; Depner *et al*., 2013b; Spooner & Jump, 2019).

Thus, while there are many studies describing different molecular and cellular effects of ω3 fatty acids (Hodson *et al*, 2017; Musa-Veloso *et al*, 2018; Okada *et al*, 2018; Šmíd *et al*, 2022; Tobin *et al*, 2018; Zöhrer *et al*, 2017) and some evidence that they may be an effective therapy for NAFLD/NASH, the key mechanisms of how they improve liver health is unknown. As such, we used a comprehensive unbiased systems approach to answer these questions. For this, we evaluated the liver transcriptome, metabolome and lipidome and assessed causal inferences via multi-omic network analysis to identify prospective mechanism operating in the diseased liver that were restored by EPA and/or DHA. Since NASH is one of the precursors of liver cancer, we also performed a meta-analysis of human liver cancer to evaluate which aspects of NASH pathogenesis leading to cancer are reversed by ω3 fatty acids. Together, our studies pointed to betacellulin (BTC), one of several epidermal growth factor receptor (EGFR) agonists, as a master regulatory molecule that was downregulated by ω3 PUFA in the NASH liver. We further validated the impact of BTC in cell culture experiments that established TGFβ-2 and integrins as the main downstream molecular targets of BTC in human hepatic stellate cells and macrophages, respectively. Suppression of these pathways specifically by DHA leads to attenuated fibrosis in NASH. Thus, our study disclosed an entirely novel mechanism for ω3 fatty acid control of hepatic function and its beneficial action against detrimental molecular events in the liver leading to NASH.

## Results

### DHA reverses the effects of WD more effectively than EPA

We first performed a comprehensive analysis of molecular changes contributing to prevention of NAFLD/NASH by two ω3 PUFA, namely docosahexaenoic (DHA, 22:6, ω3) and eicosapentaenoic acid (EPA, 20:5, ω3). For this, we evaluated transcriptomic, metabolomic and lipidomic changes caused by DHA and EPA in the whole tissue liver samples from *Ldlr ^-/-^* mice fed a western diet (WD) with the addition (or not) of DHA or EPA (Depner *et al*., 2013a) (**Figure 1A-C,** Suppl Table S1). To focus our analysis on disease features corrected by ω3 PUFA, we first established which changes induced by WD were reversed by DHA or EPA treatment (see M&M for details). We then established four categories of parameters: 1) regulated by DHA only (e.g., Cd36, **Figure 1D** top left); 2) regulated by EPA only (e.g., Notch2, **Figure 1D** top right); 3) regulated similarly by DHA and EPA (e.g., Cx3Cr1, **Figure 1D** bottom left); 4) not regulated by either DHA or EPA (e.g., Egfr, **Figure 1D** bottom right). Although there was a large overlap between the effects of each ω3 PUFA, overall, DHA showed stronger effects than EPA in restoring alterations caused by WD. Specifically, while both EPA and DHA showed similar effects on 19% of the genes affected by disease, DHA alone reversed more genes (11%) than EPA alone (3%), (**Figure 1B, C**). In line with this result, focusing on genes regulated by both fatty acids (DHA and EPA) we observed more pronounced changes by DHA than EPA (p<0.0001) (**Figure 1E, Figure S1A-B**). The genes upregulated by DHA were enriched for several functional categories: mitochondrial organization, translation, and energy derivation by oxidation of organic compounds were among the most prominent pathways affected. Regulation of cytokine production was the top enriched pathway among the downregulated genes (**Figure 1F**). Thus, the first step of transcriptome analysis indicated that DHA had a stronger effect than EPA in reversing damage inflicted by WD on the liver potentially by restoring mitochondrial function and inhibiting inflammation.

**Figure 1.**
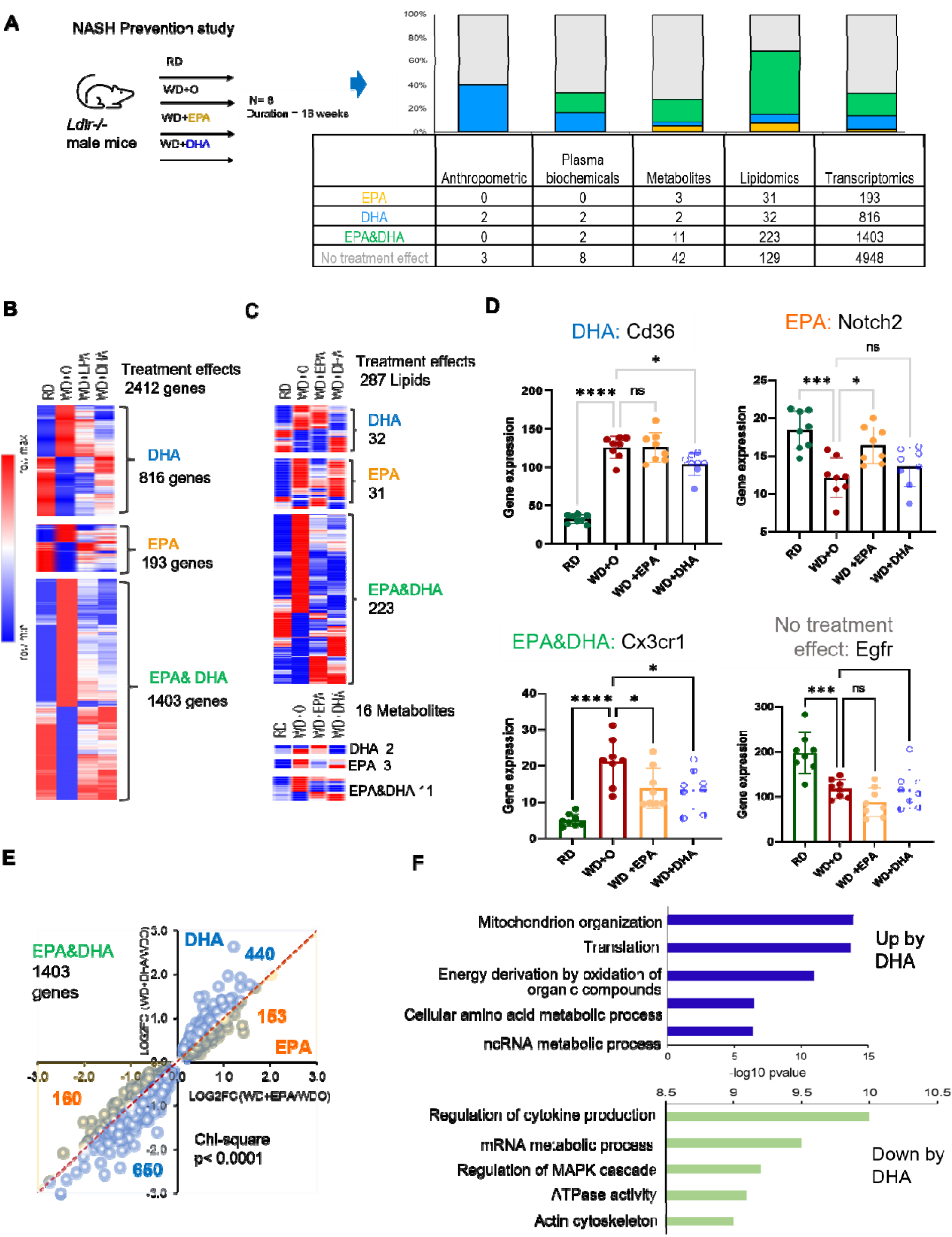
NASH mouse model outline with all expression/omics data, DHA/EPA treatment effects on outcome. A. All omics data collected and shown in Fig 1A bar graph and its associated table was acquired using samples (N = 8/treatment group) from a preclinical NASH prevention study previously described (Depner *et al*., 2013a; Depner *et al*., 2013b; Lytle *et al*, 2015). The comparison of the western diet + olive oil (WD+O) group verses reference diet (chow, RD) group showing the differentially expressed genes or parameters with a P-value <0.05 and FDR <10% and those that have treatment effect reversal uniquely by DHA (blue), EPA (orange) or similar in both EPA&DHA (green). B. Heatmap of differentially expressed genes in WD+O vs. RD fed mice and organized by treatment effect category: DHA, EPA, or DHA & EPA. Data shown are the geometric mean expression for each gene per each treatment group. Row max is displayed as red, row min is displayed as blue. C. Heatmap of parameters (lipids and metabolites) in WD+O vs. RD fed mice and organized by treatment effect category (DHA, EPA, or DHA &EPA) that are significant at P-value <0.05 and FDR <0.1. Data shown are a geometric mean of each parameter measurement for each treatment group. Row max: red, row min: blue. D. Shown are selected representatives (Cd36, Notch2, Cx3cr1 & Egfr) from each category according to treatment effect (top left: DHA, top right: EPA, bottom left: EPA & DHA, bottom right: no treatment effect). Shown here is an example of one gene from each category (Data are mean +/- SD, N=8 mice/treatment group; One-way ANOVA, with multiple comparisons test with WD+O, ns (not significant), *p<0.05, ** p<0.001, *** p<0.005, ****p<0.0001). E. Scatterplot of fold change differences between WD+O and DHA or EPA treated mice with number of genes regulated by DHA (Blue) or EPA (Orange) displayed. (Pearson’s Chi-squared test, ****p<0.0001). F. Top enriched biological process changed by DHA treatment. Top + Blue bars: Induction by DHA, Bottom + Green bars; Repression by DHA. Bars show −log10(p-value) transformed values for visualization.

In the second largest omics data set, represented by lipidomes, we did not see differences in the number of lipids regulated by DHA or EPA and ω3 PUFA restored most lipids impaired by WD (**Figures 1A, 1C)**. However, we detected a significantly stronger effect on lipids regulated by DHA versus by EPA (**Figure S1C**). Analysis of anthropometric features and plasma biochemicals also showed a more pronounced effect by DHA than EPA, as we previously reported(Depner *et al*., 2013a) (**Figure 1A**). Overall, DHA demonstrated stronger effects than EPA in reversing WD-induced changes in both gene expression and lipid concentrations.

### Mapping effects of ω3 PUFA onto multi-omic network model of NASH and its cellular components

To investigate how changes in different omics data relate to each other and which of the effects are contributing to the beneficial effects of ω3 PUFA, we have reconstructed a multi-omic network model of NAFLD/NASH and mapped ω3 PUFA effects into this model (**Figure S2A**). After filtering out data features that did not pass statistical(Dong *et al*, 2015) and causality(Yambartsev *et al*, 2016) criteria thresholds (see Materials and Methods), the multi-omic network consisted of 6743 nodes connected by 80811 edges. Specifically, the network included 6346 gene transcripts, 5 anthropometric nodes, 11 plasma biochemicals, 357 lipids, and 24 metabolites (**Figure 2A**). We next identified clusters (sub-networks) and using functional enrichment analyses found that different subnetworks were enriched in different pathways, including mitochondrial organization, myeloid leukocyte activation, cell and mitochondrial membrane fluidity, remodeling, signaling and energy metabolism. It also included processes such as macromolecule catabolic and fatty acid metabolic processes (**Figure 2A,** Suppl Table S1).

**Figure 2.**
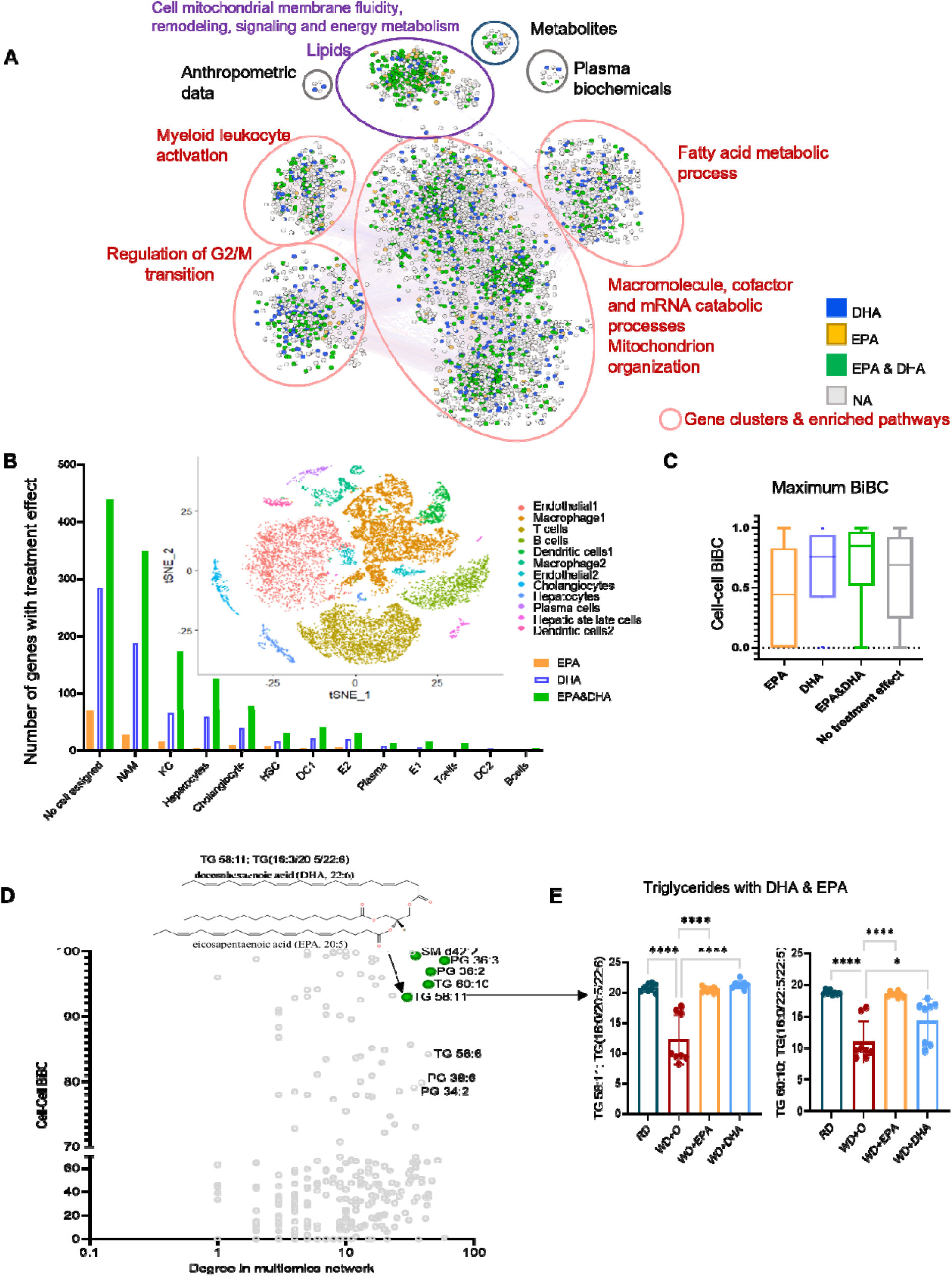
Multi-omic network (NW) reconstructed to model NASH *in vivo* using the data from preventive model (see Figure S2; M&M). A. The cytoscape visualization of the network has nodes (round rectangle) representing genes, lipids, metabolites, plasma biochemical, and anthropometric data, and edges representing correlation in a color ranging light red to light blue depending on correlation (1 to −1). The nodes are colored based on their treatment effect category membership, with DHA (blue), EPA&DHA (green), EPA (orange) and no category (grey). Network clusters are based on infomap modules additionally characterized by gene and lipids functional enrichment. B. Bar plot of number of NW genes from each treatment category (DHA (blue), EPA (orange), or EPA&DHA (green)) with assignment to a given cell type. Subplot: figure shows t-SNE plot with all cell type clusters from a reanalyzed NASH mouse single cell RNA-seq dataset (Xiong *et al*., 2019) used in our study to assign cell type information. DC- dendritic cells, HSC- hepatic stellate cells and E- endothelial cells. C. Box plot of maximum cell-cell interaction BiBC for the genes belonging to each treatment effect category (DHA (blue), EPA&DHA (green), EPA (orange) and no category (grey) shown in the graph (see Figure S2; M&M). From the network cell-cell BiBC analysis, genes regulated by DHA have higher median BiBC which indicates a higher causal contribution to cell-cell communication than genes regulated by EPA. D. The scatterplot shows the network cell-cell BiBC verses node degree for the top hepatic lipids in the NASH network. The figure insert is the structural representation of the lipid TG58:11(TG 16:0/20:5/22:6) with DHA and EPA as two of its acyl chains from the NASH prevention study. These top hepatic lipids are shown with treatment effect category EPA&DHA (green). E. Bar plots for abundance of top BiBC lipids, shown are the triglycerides with DHA and EPA as acyl chains in NASH preventive study. (Data are mean +/- SD, N=8 mice/treatment group; Ordinary One-way ANOVA, multiple comparisons test with WD+O, ns (not significant), *p<0.05, ** p<0.001, *** p<0.005, ****p<0.0001).

Multiple cell types in the liver contribute to NASH pathogenesis(Ramachandran *et al*, 2019; Seidman *et al*, 2020; Xiong *et al*, 2019) but which cells are responding to DHA treatment has not been comprehensively studied. Therefore, we mapped genes regulated by ω3 PUFA to cell type information using a previously published single cell RNA-seq dataset generated from diet-induced NASH mouse livers (Xiong *et al*., 2019). Among different cell types, most of the ω3 PUFA-regulated genes were assigned to one of the two major hepatic macrophage subpopulations including NASH-associated macrophages (NAM, 1455 genes) and Kupffer-like cells (KC, 585 genes), named according to the previous study (Xiong *et al*., 2019). These were followed by hepatocytes, cholangiocytes and hepatic stellate cells (522, 365 and 192 genes respectively) (**Figure 2B**). In line with our initial observation (**Figure 1A**), we found that DHA reversed expression of a larger number of genes than EPA, irrespectively of the cell type (**Figure 2B**).

As a next step, we wanted to ensure that our model (i.e., multi-omic network enhanced with information about cell type assignment) is consistent with known effects of ω3 PUFA on NASH. For this, we analyzed connectivity related topological network properties, known as bipartite betweenness centrality (BiBC) that was shown by us (Li *et al*, 2022; Morgun *et al*, 2015) and others (Lam *et al*, 2021) as a metric reflective of causal relationships between nodes in a co-variation network. High ranked BiBC nodes represent parameters mediating impact of one part (e.g., module/cluster) of a network on another (Li *et al*., 2022; Morgun *et al*., 2015). Thus, we calculated cell-cell interaction BiBC based on the expression of genes assigned to different cells and affected or not by ω3 PUFA in NASH multi-omic network (**Figure S2B)**. The results showed that DHA regulated genes had higher BiBCs than those regulated only by EPA or unaffected by either fatty acid (**Figure 2C; Figure S2C**). This result is in line with our previous observations of DHA being more potent than EPA in improving NASH (Lytle, 2017).

This result supported the general validity of the transcriptomic part of our network, thus, we asked which of the lipids/metabolites may have major contribution to cell-cell interactions in NASH. For this analysis, in addition to BiBC we accounted for node degree as this topological property has been the most prevalent network parameter in computational biology (Choobdar *et al*, 2019; Sorrells & Johnson, 2015) reflecting importance of the node in controlling direct and indirect neighbors and therefore corresponding biological function. Specifically, nodes of high degree (also called hubs) generally represent master regulators of a part of the large network (clusters or subnetworks) in which these hubs are situated (Choobdar *et al*., 2019; Sorrells & Johnson, 2015).

Next, we selected nodes from lipids/metabolite part of the network that were top ranked by BiBC and had high degree (top 10% for both parameters) and were also regulated by ω3 PUFA. We identified five lipids which represented less than 2% of total lipids detected in our data. Importantly, two out of the five lipids (**Figure 2D, S2D**) were triglycerides containing EPA and DHA (TG 58:11 and TG 60:10) as acyl chains (**Figure 2E**). Only 7 other lipids out of 357 had similar chemical composition (i.e., TG containing EPA/DHA) were detected by our lipidomic assay, thus making this finding being random virtually impossible (p<0.0001). Thus, this analysis identified two lipids, which are drastically depleted in the liver during and have protective effect reversing NASH (Burke *et al*., 1999; Fridén *et al*., 2021; Jump *et al*., 2015). Although this result might be expected, it is nevertheless important as a “positive control” providing additional confidence in our network.

The other three lipids were phosphatidyl glycerol (PG)36:3, PG 36:2 (cardiolipin precursors) and sphingomyelin (SM) d42:2, which are both main membrane lipids (**Figure S2D**). Since top inferences from this network validated key previously known molecular aspects of ω3 PUFA action in NASH liver we can rely on this model for inference of new yet undiscovered aspects of this process.

### Combining the NASH network with the meta-analysis of liver cancer identifies betacellulin as a key pathogenic regulator of NASH inhibited by DHA

NASH can progress to liver cirrhosis and cancer in humans (Anstee *et al*, 2019; Pfister *et al*., 2021; Samuel & Shulman, 2018). Furthermore, this was also shown in the mouse model used in this study (Chen *et al*, 2021). Although ω3 PUFA have been studied as a potential prevention strategy for colorectal cancer progression and other cancers (Dierge *et al*, 2021; Van Blarigan *et al*, 2018), there is still little understanding which cancer pathways can be inhibited by these fatty acids. Thus, as a next step, we mapped the molecular model of beneficial transcriptome alterations by DHA in liver tissue (Figure 2A) to transcriptomic alterations in human liver cancer. For this, we first performed a meta-analysis of human liver cancers (7 transcriptomic datasets with a total of 544 tumor [Hepatocellular carcinoma and Cholangiocarcinoma], 260 non-tumor, and 32 healthy patient liver samples) and established which genes were expressed concordantly by cancer and WD-induced alterations in our preclinical mouse model (**Figure S3A,** Suppl Table S2). Among 2080 concordantly expressed genes between WD and cancer, 22% (456 genes), 10% (221 genes) and 3 % (56 genes) were reversed by DHA and EPA, DHA alone and EPA alone, respectively (**Figure 3A** left panel, Suppl Table S3). While the current mouse study was designed to assess the effects of ω3 PUFA on preventing NASH (Prevention study), in another study (Lytle *et al*, 2017) we evaluated liver transcriptomes of DHA-treated mice with already established NASH (Treatment study). Preventing pathology is usually easier than to reverse it. Accordingly, we observed that treatment effects of DHA cover ∼54% (361 out of 677genes) of its preventive effects relevant for human liver cancer. Hence, these results suggest that DHA can potentially be used as cancer preventive strategy not only in patients without liver alterations, but also in those with already established NAFLD/NASH (**Figure 3A** right panel).

**Figure 3.**
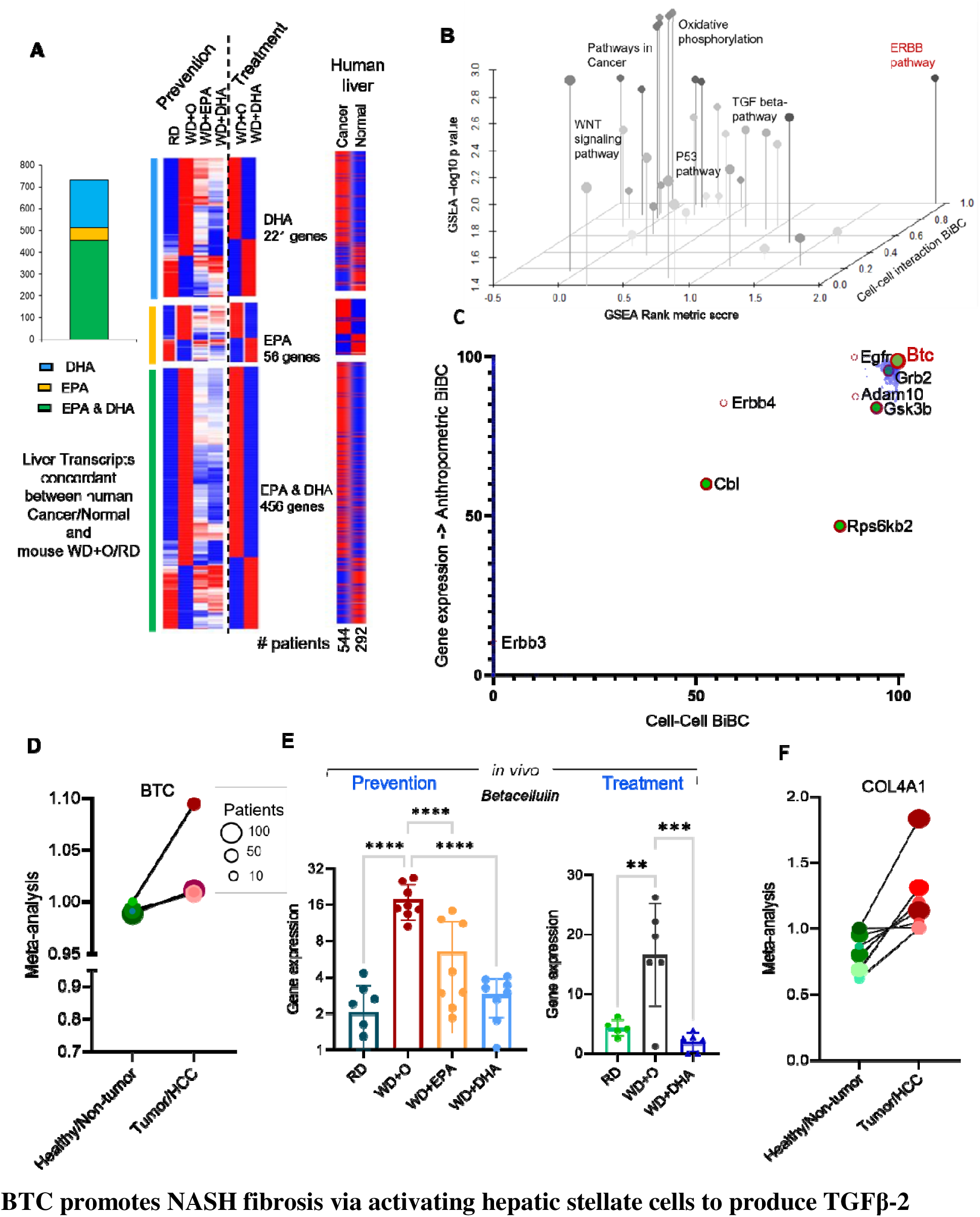
Combining NASH network with meta-analysis of liver cancer identifies betacellulin as a key pathogenic regulator downregulated by DHA. A. The heatmap for genes from human liver cancer meta-analysis (see M&M) in the right panel is organized by corresponding genes from mouse study namely, treatment effect determined by analysis of DHA or EPA effects in the prevention study and treatment study, left and center heatmaps. The stacked bar plot shows the distribution of treatment effected genes in prevention study for the genes aligned with human cancer meta-analysis genes. B. 3D scatter plot showing gene set enrichment analysis (GSEA) for the DHA effects with Rank score on the x-axis, GSEA −Log10(p-value) on the y-axis, and cell-cell interaction BiBC on the z-axis. Relevant pathways are labeled in the figure with the (BTC)-ERBB pathway (red) ranked highly by all metrics. C. Scatterplot of network BiBCs between gene expression and anthropometric parameters plotted against cell-cell BiBCs. Members of the Btc-Erbb pathway genes are overlayed and indicate the importance of Btc-Erbb signaling in the NASH multi-omic network model with preventive effects. Each circle is a node in the network, filled circles (preventive effects) with red outer circle are part of the Btc-Erbb pathway. D. BTC gene expression from meta-analysis of human normal and liver cancer datasets indicates a significant increase in liver cancer samples (Effect size FDR = 2.3 x 10^-4^). The size of each individual dot represents the number of patients associated with the dataset with higher the number of patients darker the dot color, Normal (green) and Cancer (red) (7 transcriptomic datasets with a total of 544 tumor, 260 non-tumor, and 32 healthy patient liver samples). E. Bar plot of Btc gene expression in transcriptome of mouse livers from the preventive (N=8/group) and treatment (N=5 or 6/group) experiments. (Data are mean +/- SD, Ordinary One-way ANOVA, multiple comparisons test with WD+O, ns (not significant), *p<0.05, ** p<0.001, *** p<0.005, ****p<0.0001). F. COL4A1 gene expression from meta-analysis of human normal and liver cancer data indicates a highly significant increase in expression in the liver cancer samples (Effect size FDR = 1.37 x 10^-13^). The size of each individual dot represents the number of patients associated with the dataset with higher the number of patients darker the dot color, Normal (green) and Cancer (red) (7 transcriptomic datasets with a total of 544 tumor, 260 non-tumor, and 32 healthy patient liver samples).

Next, we asked which of the molecular pathways regulated in cancer and reversed by DHA in mice may mediate the beneficial effects of DHA. For this, we combined gene enrichment analysis with BiBC to focus on genes with the largest impact on cell-cell interactions (see details in M&M). The top enriched pathway was Oxidative Phosphorylation with the well-known pathways such as TGFβ and p53 signaling being enriched to a lesser extent (**Figure S3B**). One pathway, however, that stood out as highly enriched was the ERBB signaling pathway; it was also top ranked in mediating DHA-driven cell-cell interactions (i.e., BiBC) (**Figure 3B, Figure S3C**). ERBBs are known homologs of EGFR, which are activated through binding to EGF and related members of the EGF family of growth factors. These include EGF-like ligands or cytokines that are comprised of at least ten proteins including betacellulin, transforming growth factor-alpha (TGF-α), amphiregulin, HB-EGF, epiregulin, and neuregulins and the various other heregulins (Chen *et al*, 2016; Olayioye *et al*, 2000; Wieduwilt & Moasser, 2008).

Using our multi-omic network (including DHA affected and not affected genes in NASH) we ranked genes from the ERBB pathway based on their potential capacity to mediate effects of DHA and identified betacellulin (BTC), an alternative ligand of EGFR (Figure S2D), as a top gene among the important genes (Grb2, Gsk3 Gsk3β/α and Cbl) in the downstream pathway (**Figure 3C**). Interestingly, EGFR itself, although not regulated by DHA was the second best potential regulatory gene for this pathway in NASH. In contrast, BTC was found to be increased in liver cancer meta-analysis (**Figure 3D**), up regulated by WD and downregulated by DHA both in prevention and treatment mouse studies (**Figure 3E**). To ensure the statistical robustness of BTC’s potential causal capacity reflected by its high BiBC rankings, we used two additional approaches (non-parametric statistics that would identify outlier values of BiBC and “experimental” statistics by comparing a given node BiBC value to simulated random networks). Each approach demonstrated extremely low probability of BTC being randomly highly ranked (with p<10^-15^ and probability density =0.009, respectively) (**Figure S3D-E**). Thus, we established BTC as a top causal candidate gene mediating DHA’s beneficial effect on hepatic health, while also being relevant for liver cancer in humans.

Given that BTC was predicted as a new target, central for effects of DHA in NASH, we next verified which cell types express BTC and its receptor (EGFR) in our model of NASH. For this, we integrated the network with available interaction information from ligand-receptor database (Abugessaisa *et al*, 2021) and single cell RNA-seq data (Xiong *et al*., 2019) (**Figure S3F**). First, we observed that cholangiocytes were the primary population of cells expressing BTC (**Figure S4A**). Although its source is restricted to cholangiocytes, BTC’s role as a secreted growth factor indicates it can act on many different types of neighboring cells (e.g., hepatocytes, macrophages, hepatic stellate cells and others) that express EGFR/ERBBs (**Figure S3F, Figure S4B**).

Among different liver cell types which can respond to BTC and proliferate during NASH progression, hepatic stellate cells (HSCs), frequently called mesenchymal cells in humans (Carter & Friedman, 2022; Ramachandran *et al*., 2019) produce collagens and represent a major contributor to hepatic fibrosis. Indeed, the number of mesenchymal cells counted in humans with cirrhosis is markedly higher than those of healthy livers (**Figure S4C**). Importantly, DHA prevents and reverses fibrosis in a NASH mouse model (Depner *et al*., 2013a; Lytle *et al*., 2015) and decreases expression of two out of three collagen encoding genes (COL1A1, COL1A2, COL4A1) that are upregulated in liver cancer (**Figure 3F, S4D,** Suppl Table S3). Therefore, we next tested effects of BTC on human hepatic stellate cells using the LX2 cell line (Xu *et al*, 2005). LX2 cells were grown and pretreated with and without BTC using EGF as a growth factor positive control. LX2 growth was significantly increased by BTC (**Figure 4A**) and to a similar extent as was observed for EGF (**Figure S4E**). Moreover, we observed increased collagen staining (Sirius red) in cells treated with BTC (**Figure 4B**).

**Figure 4.**
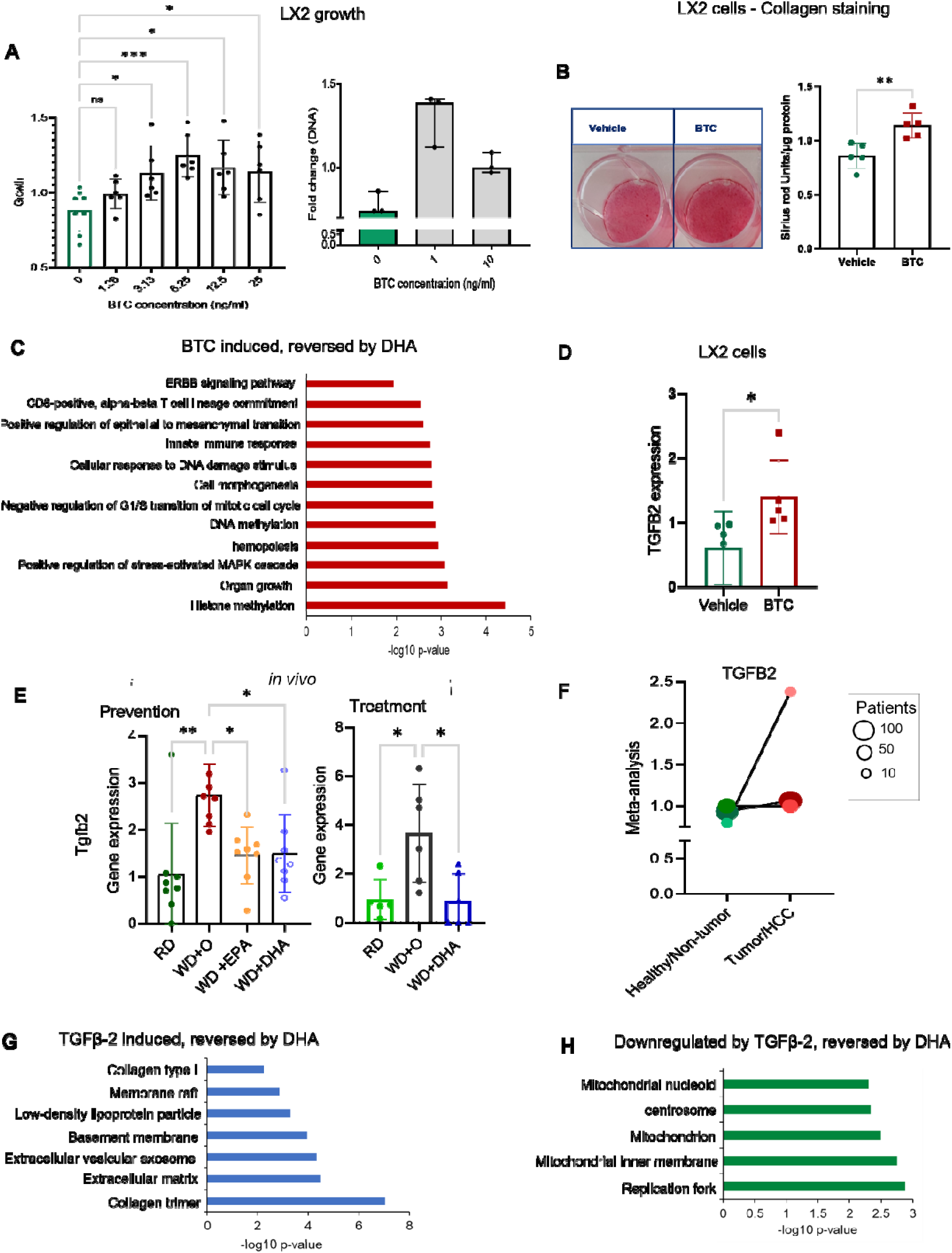
Hepatic stellate cell (LX2) proliferation could promote fibrosis via BTC-TGFβ-2, reversed by DHA. A. LX2 cell proliferation assay after treatment with BTC, 0-25 ng/mL (left panel, N=6 individual experiments). Fold change in DNA concentration after treatment with BTC 0-10 ng/mL (right panel, N=3 experiments). Green bar in each plot is untreated control. Data are displayed as mean +/- SD with each point being an individual experiment; Ordinary One-way ANOVA, multiple comparisons test with control vehicle, ns (not significant), *p<0.05, ** p<0.001, *** p<0.005, ****p<0.0001). B. Collagen staining (Pico Sirius Red, PSR) in LX2 cells indicating increased fibrosis (collagen production) when stimulated with BTC (20ng/ml; N=5 experiments) or a vehicle control. A representative image of stained cells is shown (left panel). A bar plot of staining intensity normalized by total protein (µg) per well. (N=5 experiments; unpaired, two-sided t-test **p=0.0058). C. Gene enrichment (biological process) of LX2 cells (see M&M) while induced by BTC (BTC vs Vehicle; one-sided t-test) identifies pathways reversed by DHA treatment *in vivo.* Data are displayed as −log10(p-value). D. Bar plot of TGFB2 gene expression in LX2 cells treated with vehicle or BTC (20 ng/ml; N=5 experiments) *in vitro* (paired, two-sided t-test *p<0.05, FDR<0.1). E. Tgfb2 gene expression *in vivo.* DHA reversed the gene expression significantly in the *in vivo* experimental model both in Preventive & Treatment models. (Data are displayed as mean +/- SD, N=8 mice/group (preventative study) or N=5 or 6 mice per group (treatment study); Ordinary One-way ANOVA, multiple comparisons test with control vehicle, ns (not significant), *p<0.05, ** p<0.001, *** p<0.005, ****p<0.0001). F. TGFB2 gene expression from meta-analysis of human normal and liver cancer data indicates the significant increase in expression in cancer samples (Effect size FDR <0.0064). The size of each individual dot represents the number of patients associated with the dataset with higher the number of patients darker the dot color, Normal (green) and Cancer (red) (7 transcriptomic datasets with a total of 544 tumor, 260 non-tumor, and 32 healthy patient liver samples). G. Gene enrichment analysis of genes upregulated in TGFβ-2 treated cells (GSE45382) that were reversed by DHA *in vivo,* identifies top enriched pathways. Data are displayed as −log10(p-value). H. Gene enrichment analysis of downregulated genes in TGFβ-2 treated cells (GSE45382) that were reversed by DHA *in vivo,* identifies highly enriched gene ontologies. Data are displayed as −log10(p-value).

We next performed a RNASeq transcriptomic analysis of LX2 cells treated with BTC and compared it to genes regulated by DHA in vivo. We found 63 genes upregulated by BTC and downregulated by DHA and 16 genes downregulated by BTC and upregulated by DHA. Among the enriched pathways for the set of genes induced by BTC and repressed by DHA were transcripts involved in cell growth including ERBB signaling (**Figure 4C, S4F-H,** Suppl Table S4). Strikingly, TGFB2 was the only gene found in common across several enriched categories. Moreover, expression of TGFB2, but not TGFB1 was increased by BTC in vitro (**Figure 4D, S4G),** repressed by DHA in both prevention and treatment mouse studies (**Figure 4E),** and increased in human liver cancer meta-analysis (**Figure 4F**). A recent study demonstrated that TGFβ-2, but not TGFβ-1 has a critical non-redundant role in promoting lung and liver fibrosis (Sun *et al*, 2021). Therefore, we hypothesized that downstream effects of reduction of BTC by DHA can be explained by reduction of TGFβ-2. For this, using publicly available *in vitro* data on TGFβ-2 effects and our *in vivo* data, we evaluated which genes regulated by DHA were regulated in the opposite direction by TGFβ-2 (Suppl Table S5). We found 62 genes upregulated by TGFβ-2 and downregulated by DHA and another 62 genes downregulated by TGFβ-2 and upregulated by DHA. DHA downregulated/ TGFβ-2-upregulated genes were highly enriched for production of collagen trimers and extracellular matrix organization (**Figure 4G**) while genes downregulated by TGFβ-2 and reversed by DHA were enriched for mitochondrial inner membrane and other mitochondrially related functions (**Figure 4H**). Altogether these results suggest that one of a key mechanism of fibrosis reduction by DHA is achieved through an inhibition of BTC and consequent reduction of HSCs proliferation and TGFβ-2-induced collagen production (Sun *et al*., 2021).

### TLR-dependent inflammatory processes in NASH are exacerbated by BTC

Our results from stellate cells support that inhibition of BTC by DHA would explain reduction of fibrosis and improvement of mitochondrial function. However, the reduction in inflammatory pathways and macrophage gene expression, another major effect of DHA we observed (Fig. 1 and 2), could not be explained by effects of BTC on stellate cells. Furthermore, EGFR deficiency specifically in macrophages has been shown to attenuate liver cancer in a mouse model (Lanaya *et al*, 2014b). Finally, we have previously reported that systemic levels of TLR2 ligands are decreased by DHA (Lytle *et al*., 2015) (**Figure S5A**). These results, along with a well-known fact that a genetic deficiency of microbiota sensors (TLR2 and TLR4) attenuates NASH (Miura *et al*, 2013; Spruss *et al*, 2009; Wu *et al*, 2020), indicate that a reduction in TLR2 and/or TLR4 ligand levels by DHA might interact with the reduction of BTC in preventing the disease. While different receptors for BTC are widely distributed across different liver cell types (Figure S4B**, S5B**), TLR2 and TLR4 are predominantly expressed by macrophages in the liver (**Figure S5C**).

Taken together, our next question was: which processes regulated by DHA in the liver can be explained by the effects of BTC and TLR2/4 agonists on macrophages? We also asked whether BTC modulates TLR2/4-dependent immune stimulation in macrophages. To answer these questions, we differentiated the human monocyte cell line (THP-1) to a macrophage-like phenotype and stimulated with BTC, LPS (TLR4 ligand) and PGN (TLR2 ligand) and compared global gene expression in these cells to cells stimulated with TLR2/4 ligands only or unstimulated control cells (**Figure S5D,** Suppl Table S6).

To investigate a potential interaction effect, we tested a range of concentrations of BTC and TLR-agonists evaluating expression of IL-8, and CCL2 (MCP-1) expression, well-known targets of BTC (Lanaya *et al*., 2014b; Shi *et al*, 2014) and TLR-agonists (Miura *et al*., 2013; Seki *et al*, 2007; Spruss *et al*., 2009; Wu *et al*., 2020) and chose the lowest doses that induces their expression (**Figure S5E, S5F**). We next performed transcriptomic analysis of THP-1 cells treated with BTC-LPS-PGN and compared it to genes regulated by DHA *in vivo*. We found 179 genes upregulated by BTC-LPS-PGN and downregulated by DHA and 285 genes downregulated by BTC-LPS-PGN and upregulated by DHA (**Figure 5A)**. Genes upregulated by BTC-LPS-PGN were enriched for several pathways related to monocyte/macrophage related immune functions, the cell cycle, and collagen binding. Among the most enriched categories for the downregulated genes were mitochondrion, NAD metabolic process, and endoplasmic reticulum membrane (**Figure 5B-G)**.

**Figure 5.**
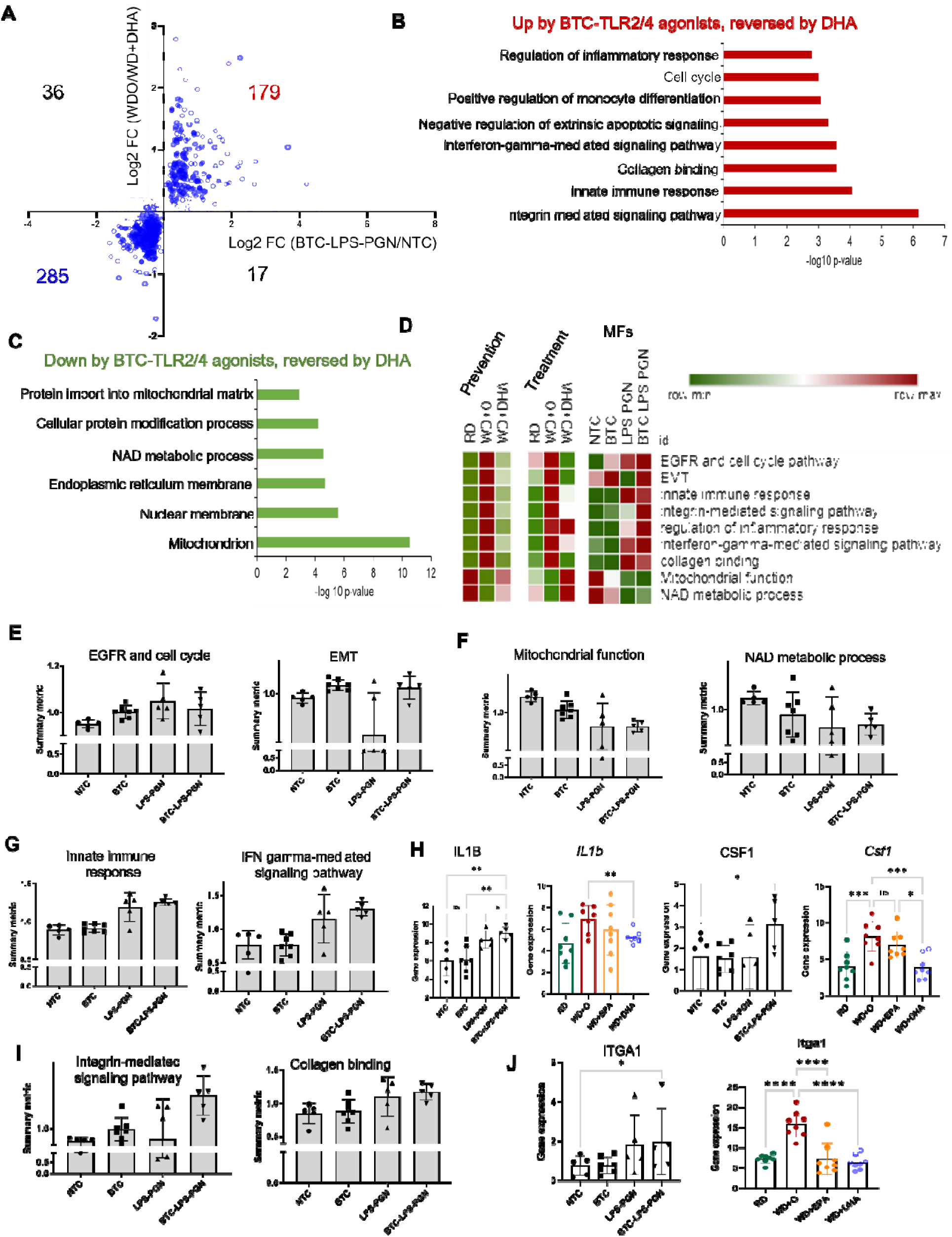
BTC promotes TLR-dependent inflammation and integrin production by macrophage (THP1) cells. A. Scatterplot of differential expression with BTC-LPS-PGN (BTC and TLR2/4 ligands) treated cells to control THP-1 cells *in vitro* (BTC-LPS-PGN/control in THP-1 cells; p-value <0.05; Details about concentrations and duration of the TLR agonists treatment are detailed in M&M) against *in vivo* differential expression with WDO to WD+DHA in NASH preventive model. Filled circles are the genes in DHA treatment category. B. Gene enrichment is shown for the genes upregulated from BTC-LPS-PGN treatment in THP-1 cells that were reversed by DHA treatment *in vivo*. (Gene ontology-biological process. Data are displayed as −log10(p-value). C. Gene enrichment is shown for the genes downregulated from BTC-LPS-PGN treatment in THP-1 cells that were reversed by DHA treatment *in vivo*. (Gene ontology-biological process and cellular components). Data are displayed as −log10(p-value). D. The heatmap for summary metric of all major molecular pathways affected by BTC treatment in THP- 1 cells shown to be prevented by DHA in both *in vivo* NASH preventive and treatment models (see M&M). E-G. The individual pathways enriched in BTC-LPS-PGN treated THP-1 cells but reversed by DHA *in vivo* in the NASH preventive and treatment models displayed as summary metric. H. Individual gene expression for IL-1b and CSF1 from THP-1 cells *in vitro* treated with BTC and or TLR2/4 ligands (N=5 experiments, paired, one-sided t-test, ns (not significant), *p<0.05, ** p<0.001) (see M&M) and *in vivo* NASH preventive model (Data are mean +/- SD, N=8 mice/treatment group). (Ordinary One-way ANOVA, with Dunnett’s multiple comparisons test with WD+O, ns (not significant), *p<0.05, ** p<0.001, *** p<0.005, ****p<0.0001). I. The bar plots for summary metric are shown for integrin signaling and collagen pathway in THP1 cells. J. The bar plot for ITGA1 gene expression is shown in THP-1 cells *in vitro* treated with BTC and or TLR2/4 ligands (N=5 experiments, paired, one-sided t-test, ns (not significant), *p<0.05, ** p<0.001) (see M&M) and *in vivo* NASH preventive model (Data are mean +/- SD, N=8 mice/treatment group). (Ordinary One-way ANOVA, with Dunnett’s multiple comparisons test with WD+O, *p<0.05, ****p<0.0001).

To assess the relative contribution of BTC and of TLR2/4 agonists to the observed combined functional effect, we calculated a summary metric for each pathway (see M&M) and compared their values between each treatment group (**Figure 5D)**.

To check if the expected effects of BTC are present in macrophages, we first verified expression of EGFR/cell cycle and epithelial-mesenchymal transition (EMT) pathways and observed their increase among the categories upregulated by BTC alone and in combination with TLR agonists (**Figure 5E)**. Analysis of the downregulated pathways showed that in combination with TLR ligands BTC inhibited expression of genes involved in mitochondrial functions (TCA cycle, oxidative phosphorylation, **Figure S5G**) and NAD metabolic functions (critical pathways operating in mitochondria) (Samuel & Shulman, 2018; Simões *et al*, 2018; Xie *et al*, 2020) (**Figure 5F**).

As for the upregulated pathways, we observed diverse immune functions such as ‘innate immune response’, ‘regulation of interferon-gamma signaling’ with a similar or slightly enhanced expression of genes when BTC was added with TLR2/4 agonists (**Figure 5G, Figure S5H**). However, some genes showed clear interactions between BTC and TLR-agonists. For example, IL1B was upregulated by TLR2/4 agonists and slightly increased by BTC, but CSF1, a classical factor of macrophage growth and proliferation (Hume & MacDonald, 2012), was upregulated only when both BTC and TLR2/4- agonists were present (**Figure 5H**).

Among immune related pathways, the strongest BTC effect either alone or in combination with TLR2/4 agonists was on the integrin-mediated signaling pathway, which partially overlapped with collagen binding (**Figure 5I**). Interestingly, while there was a trend for BTC increasing transcript levels of ITGα6 and ITGα9 and TLR2/4 agonists of ITGα1, only the combination of BTC with TLR-agonists significantly induced expression of all three integrins (**Figure 5J, S5I**). Notably, the integrin pathway was not regulated by TLR2/4 agonists alone suggesting a possible unique role of the crosstalk between EGFR and TLR pathways in controlling fibrosis related molecular function in macrophages **(Figure 6).**

**Figure 6.**
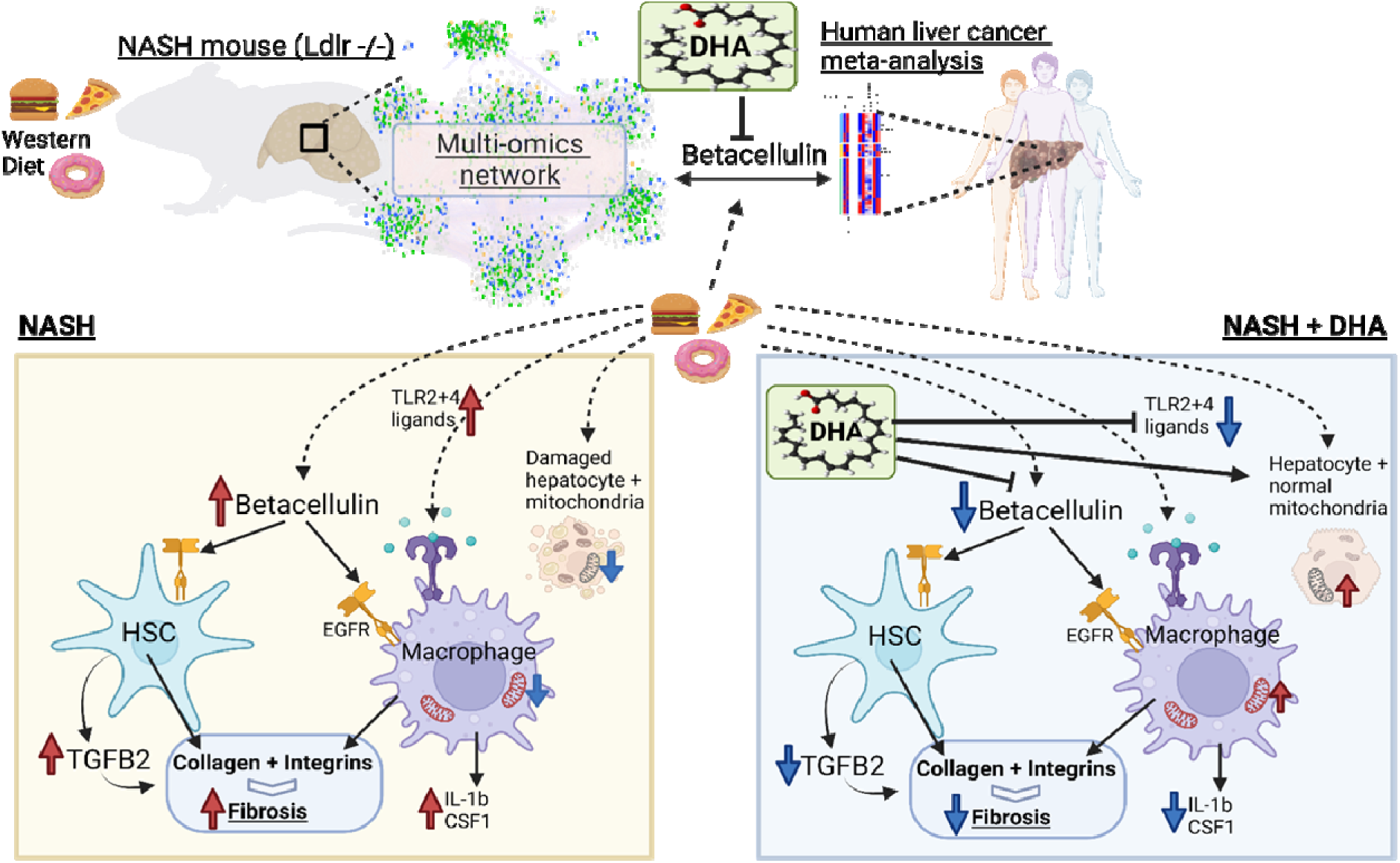
Proposed mechanism of attenuation of NASH by DHA. Based on mouse and human single cell RNASeq studies, BTC is expressed in cholangiocytes and acts locally to regulate the ERBB pathway in various liver cells, e.g., hepatic stellate cells (HSC) and macrophages. NASH is associated with increased BTC expression in human and mouse livers. DHA attenuates BTC expression in livers of mice fed the western diet. BTC effects on LX2 cells (HSC) include increased cell proliferation and TGFβ-2 expression leading to collagen production, while BTC effects on THP-1 cells involves increased production of key cytokines and integrins. DHA prevents these pathways by lowering the BTC’s action on these cells leading to decreased fibrosis, hepatic inflammation and improving mitochondrial function as seen *in vivo*.

## Discussion

For many metabolic diseases such as diabetes (Dahlén *et al*, 2021; Jermendy *et al*, 2018), atherosclerosis (Bhatt *et al*, 2019), and obesity (Jastreboff *et al*, 2022) there are highly efficient drugs that treat and/or prevent development of these diseases. The only frequent representative of metabolic diseases that still lacks pharmacological agents that would pass clinical trials is NAFLD/NASH. As this disease is often called “fatty liver”, most of attempted treatments target reduction of liver fat (Rinella, 2015). However, these treatments cannot resolve liver fibrosis, which is a more resistant aspect of the pathogenesis of this disease (Wattacheril *et al*, 2018). Fibrosis is the main cause of liver failure in patients with NASH and also precedes and leads to development of liver cancer (Anstee *et al*., 2019; Pfister *et al*., 2021). Therefore, revealing the mechanisms of action of DHA that reduce fibrosis in a preclinical mouse model may aid in the rational use of ω3 PUFA or help in developing new drugs that would act upon the same cellular/molecular targets (Jump *et al*., 2015).

In this study, we identified betacellulin (BTC), one of the less studied ligands of EGFR, as a master regulator whose reduction by DHA potentially leads to prevention/treatment of fibrosis (**Figure 6**). Indeed, we revealed that BTC induces the TGFβ-2, a critical contributor to liver fibrosis via collagen production by hepatic stellate cells (Dropmann *et al*, 2016). Moreover, in combination with TLR 2/4-agonists (also reduced by DHA), BTC induces integrin pathway in macrophages, the cell type in the liver most affected by DHA treatment and well-known to be involved in pathogenesis of fibrosis in different organs (Wynn & Vannella, 2016). Thus, BTC represents a candidate master regulator inducing two most important factors (collagens and integrins) contributing to liver fibrosis and consequently promoting liver cancer.

In addition to its effect on fibrosis, reduction of BTC seems to be also mediating another important effect of DHA, which is improvement of mitochondrial function related pathways. Indeed, mitochondrial damage is widely reported in NAFLD/NASH and its improvement by DHA is clearly seen even at the initial stage of our analysis (Figure 1E) indicating a strong impact of DHA on this pathway. Accordingly, BTC in combination with microbiota derived stimulation (represented by TLR agonists) has a negative effect on the expression of genes involved in mitochondrial functions.

Another robust effect of DHA observed at the initial step of our analysis was the inhibition of inflammation. Not surprisingly, a deeper investigation into this phenomenon led us to potential effects of DHA on microbiota-related molecules and on hepatic macrophages involved in sensing microbes. In fact, network analysis combined with scRNAseq data pointed to macrophages as main cellular targets of DHA in the liver (Figure 2B). Macrophages are also the primary cells in the liver that express both TLR 2/4 (Figure S5C) whose microbiota-derived agonists are decreased by DHA (Figure S5B) (Lytle *et al*., 2015). This was an important observation considering that deficiency of either TLR2 or TLR4 attenuates severity of NADFL/NASH in mouse models (Miura *et al*., 2013; Spruss *et al*., 2009; Wu *et al*., 2020).

In contrast to TLRs, EGFR and other receptors that BTC binds are widely expressed across most cells in liver including macrophages. Moreover, EGFR deficiency in macrophages, but not in hepatocytes was shown to attenuate NASH in mice (Lanaya *et al*, 2014a)

Thus, it is plausible that the impact of DHA on macrophage related to NAFLD/NASH pathogenesis can be explained by the fact that DHA simultaneously limits cell access to BTC and TLR2/4 agonists. Moreover, despite the limitations of our cell culture system and the difficulty of transition from mice to humans, we observed that BTC combined with TLR 2/4 agonists induce the integrin signaling pathway which was inhibited by DHA in the liver. Inspecting individual genes revealed that all detected integrins (ITGα1,6,9- Figure 5J, Figure S5I) required all three compounds for induction except for ITGβ1 that increased only with only BTC stimulation in THP-1 cells. Interestingly, ITGβ1 was also increased by BTC in hepatic stellate cells (Figure S4H). Of note, proteins coded to ITGαs and ITGβs form a complex that binds collagens (Bourgot *et al*, 2020). Accordingly, blocking integrin signals has been shown to attenuate fibrosis (Agarwal, 2014; Rahman *et al*, 2022). Hence, our results taken together with already existing literature about other pathologies (Bourgot *et al*., 2020) suggest that DHA inhibits the macrophage contribution to fibrosis by simultaneously inhibiting microbiota-derived signals and BTC. Furthermore, while BTC is a new player in this arena, the role of microbiota derived signals in activating a profibrotic program in macrophages has been reported for different diseases in a few organs (Costa *et al*, 2022; He *et al*, 2021).

Leveraging complex multi-omic and single cell data (Xiong *et al*., 2019) and using a systems approach, our study constructed a statistical network model of cell-cell interactions affected by DHA (Figure S3F). More importantly, network cause-effect related information flow (degree and BiBC) combined with a ligand-receptor database(Abugessaisa *et al*., 2021) (Figure S3F) allowed us to infer that a candidate master regulator molecule (BTC) produced by one cell (cholangiocytes) acts upon several other cells. DHA may affect several cell types involved in liver fibrosis. Our study, however, reveals that BTC inhibition by DHA simultaneously disrupts the integrin pathway in macrophages and TGFβ-2 -driven collagen production by hepatic stellate cells, two processes that synergize in the development of liver fibrosis. Thus, we propose that removal of BTC and Tlr2/4 agonists prevents binding of integrins to collagen that is required for the scar development.

The main outcomes of our study, however, might miss some additional beneficial effects of DHA, especially those that are not related to BTC reduction and fibrosis. This is because we focused our investigations on cellular/molecular events relevant to NASH and its progression to liver cancer in humans. Specifically, while our network analysis modeled NASH in mice, we used it as a first step in identification of the most critical causal pathways upregulated in hepatic cancer meta-analysis and reversed by DHA. In this analysis we found ERBB as a top pathway with BTC being the top-ranked molecule in this pathway altered by DHA. Accordingly, our *in vitro* investigations of human hepatic stellate cells demonstrated that BTC promotes cell growth, which was an expected finding considering that BTC belongs to a family of growth factors (Chen *et al*., 2016; Olayioye *et al*., 2000; Wieduwilt & Moasser, 2008). Thus, in addition to promoting fibrosis by upregulation of TGFB2 in stellate cells, it may also increase numbers of these collagen-producing cells in the liver. Furthermore, our gene expression analyses of effects promoted by BTC *in vitro* and downregulated by DHA in the liver also support this growth function for BTC (Figure 4A), pointing to an activation of ERBB pathways, cell growth and even the epithelial mesenchymal transition. Dysregulation of these pathways are hallmarks of cancer (Chava *et al*, 2022; Dahlhoff *et al*, 2014; Lanaya *et al*., 2014b). The growth-promoting actions of BTC are known to be mediated by epidermal growth factor receptors (ERBBs), namely ERBB1(EGFR) (Chava *et al*., 2022), ERBB2, ERBB3 and ERBB4. In liver however, the mechanism for BTC dependent cell proliferation has not been elucidated. The BTC-EGFR- ERBB4 pathway in pancreatic ductal adenocarcinoma has been well established by several groups and a BTC knock out can partially rescue the cancer progression (Hedegger *et al*, 2020). In the *in vivo* NASH model, we have a significant reversal of ERBB4 expression by DHA (Figure S5B). Furthermore, DHA also inhibits GRB2 and GSK3β, genes which are downstream in the ERBB signaling pathway found in our liver cancer meta-analysis (Figure 3A, C) and known to promote multiple types of cancers (He *et al*., 2021). These results suggests that DHA, besides its main effect as BTC reduction, might also inhibit residual activation of ERBB pathway via BTC-unrelated mechanisms. In addition, forty-nine EMT related genes, including the TGFβ- 2 mediated (Dropmann *et al*., 2016) SMAD-EMT-cancer pathway were reversed by DHA treatments (Figure 3A). Therefore, our results suggest that the DHA attenuation of BTC-TGFβ-2-dependent molecular events might not be limited to reversal of fibrosis in NASH (Lytle *et al*., 2015) but also has a promise in preventing NASH’s progression into liver cancer. Despite the novelty and importance of our findings several questions remain to be investigated. For example, to evaluate the relative contribution of down regulation of BTC by DHA for treatment of NASH and prevention of cancer, experiments using mouse models might be needed.

Clinical trials using EPA and/or DHA in NAFLD/NASH therapy have shown mixed, but promising results with significant improvement with disease severity, including hepatosteatosis and liver injury (Hodson *et al*., 2017; Musa-Veloso *et al*., 2018; Okada *et al*., 2018; Šmíd *et al*., 2022; Tobin *et al*., 2018; Zöhrer *et al*., 2017). Unfortunately, these studies did not address whether patients that did not respond to therapy represent a cellular/molecular subtype of NAFLD that is not treatable only by ω3 PUFA or perhaps have such severe disease that they are unlikely to respond to any therapeutic agent.

In conclusion, with the discovery of BTC as a candidate to be one of the key mediators of ω3 PUFA therapeutic effects, our study opens a new avenue for investigation of NAFLD/NASH. In addition to finding new mechanisms of action of DHA, this study is the first to demonstrate that BTC can induce TGFβ-2 and synergize with microbial signals in the induction of integrins. Thus, while few earlier studies (Moon *et al*, 2006) showed increase of BTC in the liver tumors, our robust meta-analysis coupled with evidence for causal contributions shed a new light to this molecule in the pathogenesis of this cancer. Moreover, BTC’s role in human NAFLD/NASH is entirely uncharted territory. Therefore, future studies should investigate if BTC-triggered gene expression signatures can serve as biomarkers guiding personalized ω3 PUFA therapy, as targets of new NAFLD/NASH drugs, and finally as a predictors of hepatic cancer risk in humans.

## Acknowledgments

We would like to thank Dr. Scott Friedman from the Division of Liver Diseases at the Icahn School of Medicine at Mount Sinai for providing the LX2 cell line and the personnel of CQLS at the Oregon State University for IT support.

## Author contributions

Conceptualization: JP, NKN, MGJ, JWP, AM, NS, DBJ Methodology: JP, NKN, SS, PM, MGJ, JWP, RR, ZL, AKD, KB, AM, NS, DBJ Investigation: JP, NKN, MGJ, JWP, PM, RR, ZL, AKD, KB, GT, AM, NS, DBJ Visualization: JP, NKN, JWP, AM, NS, DBJ Funding acquisition: AM, NS, DBJ, GT Project administration: NS, DBJ, AM Supervision: AM, NS, DBJ Writing – original draft: JP, NKN, NS, AM, DBJ Writing – review & editing: JP, NKN, MGJ, JWP, SS, RR, ZL, AKD, GT, KB, NS, DBJ, AM

## Data and materials availability

RNA-Seq data: Gene Expression Omnibus GSE215223, GSE215224, GSE215225 and GSE215227.

The complete metabolomic dataset can be here accessed https://www.ebi.ac.uk/metabolights/MTBLS6001.

## Supplementary Materials

**Fig. S1** to **Fig. S6** for multiple supplementary figures with the legends.

**Table S1.** The table shows all differentially expressed genes (DEGs), parameters between WD and ND in mice (FDR<10%) and DEGs that were retained in the network from the liver NASH prevention and treatment study model and their cell type assignments.

**Table S2.** Human liver cancer datasets and sample details used in meta-analysis.

**Table S3.** Meta-analysis of human liver cancer data.

**Table S4.** Gene expression from LX2 cells with or without BTC treatment mapped on to the NASH network genes.

**Table S5.** Gene expression from TGFβ-2 treatment mapped on to NASH network genes.

**Table S6.** Gene expression from THP-1 cells with or without BTC and LPS/PGN treatment mapped on to the NASH network genes.

## Supplementary Figures

**Suppl Figure 1.**
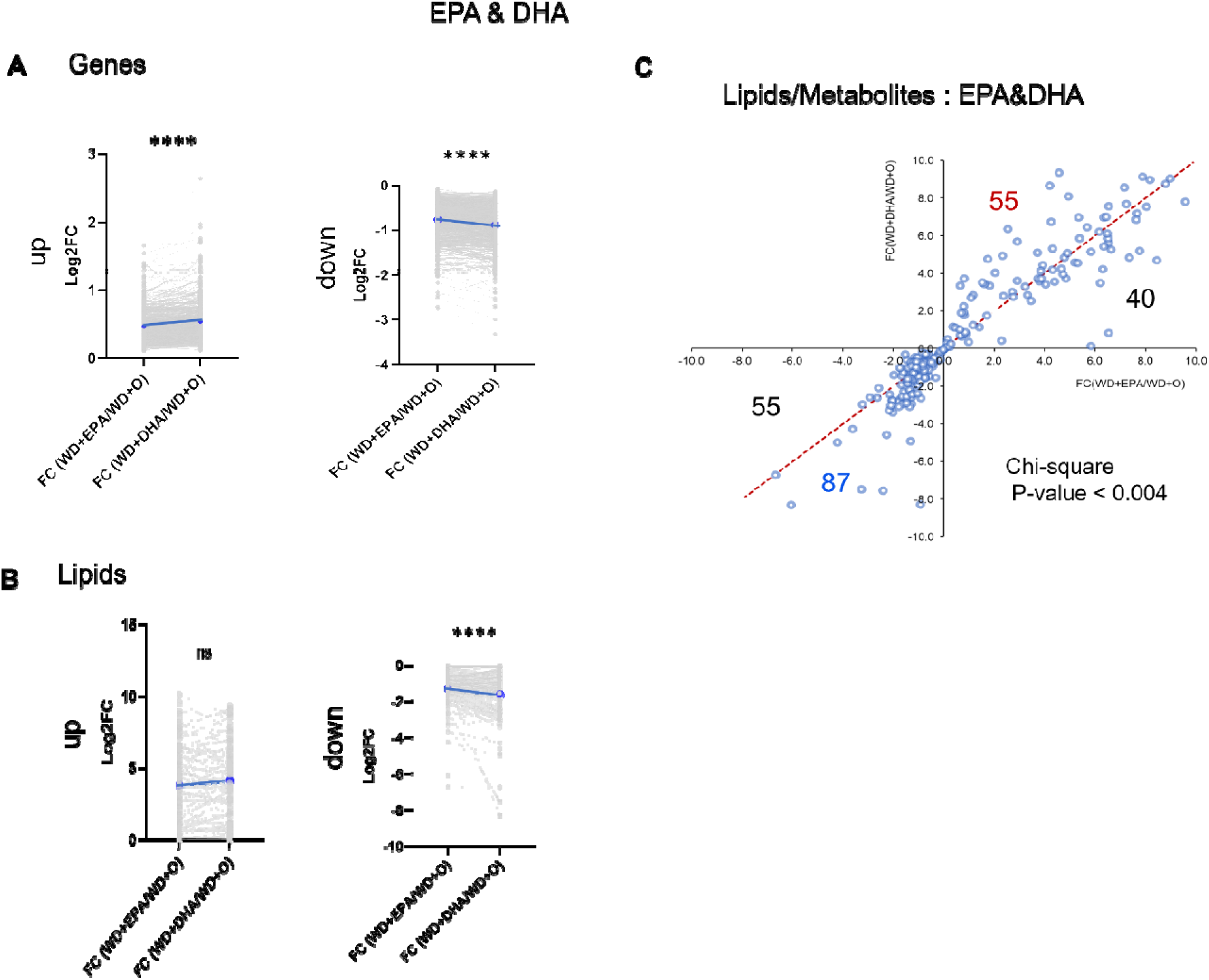
A-C. The Log2FC of EPA & DHA gene expression (A) and lipids and metabolites (B-C) show the extent of DHA reversal effects as significantly higher than EPA though similar in profile. The up and down regulation are shown in separate plots for clarity (A-B) (paired, two-sided t-test, ns (not significant), ****p<0.0001). C. Scatterplot of fold change differences between WD+O and EPA (x-axis) or DHA (y-axis) treated mice with number of lipids and metabolites regulated similarly by DHA & EPA displayed. (Pearson’s Chi-squared test, **p<0.004).

**Suppl Figure 2.**
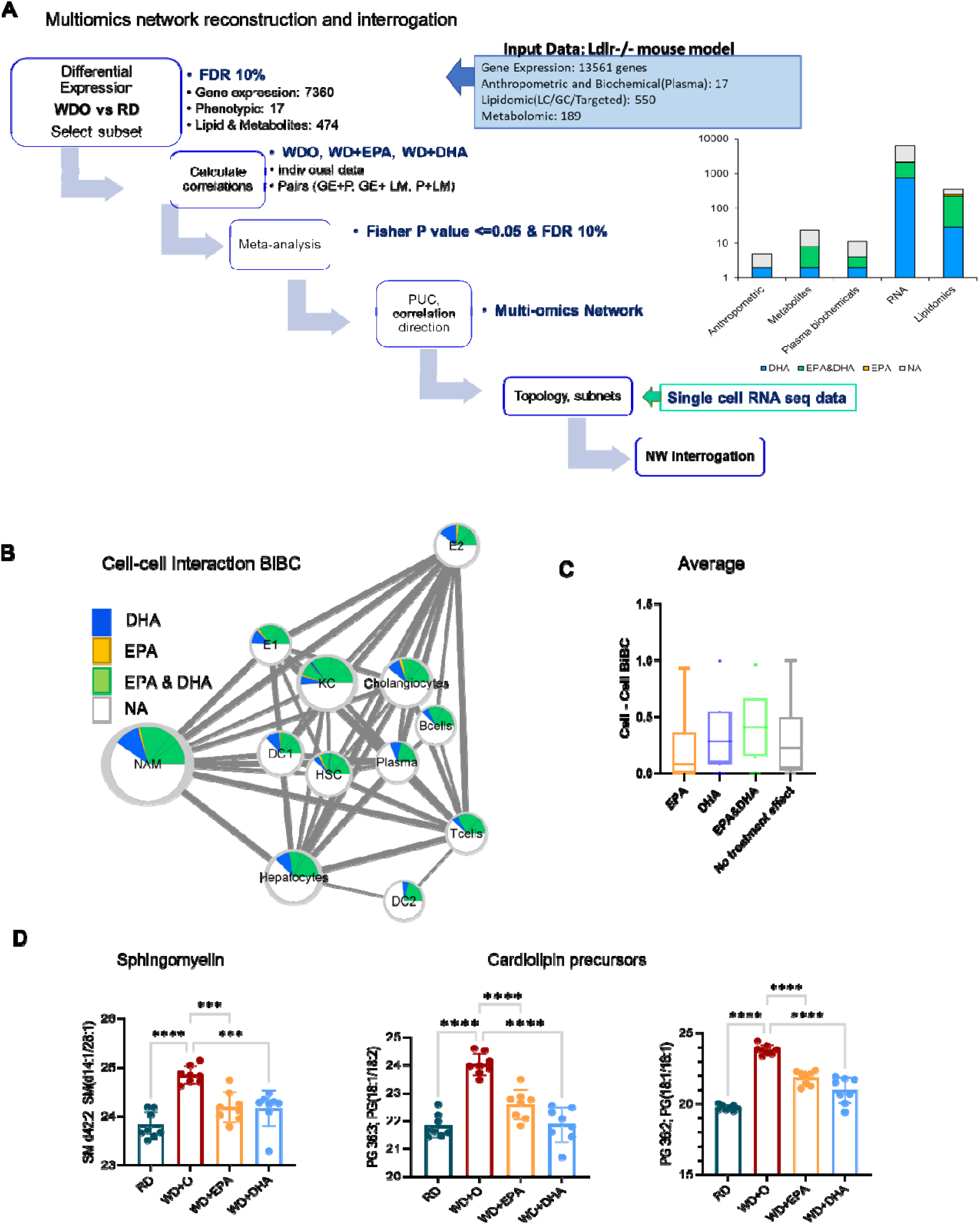
A. An outline with the steps involved in deriving, analyzing and interrogation of network model from the multi-omic NASH data (see M&M) (GE: gene expression; P: Phenotype/Anthropometric/Biochemical data; LM: Lipids/Metabolites; PUC: Proportion of unexpected correlation). The distribution of treatment effects from the NASH preventive model among the network parameters is shown in the right panel. B. Outline of cell-cell interactions used to calculate the interaction BiBCs (see M&M). The nodes represent all genes part of cell types as a cluster in the network and edges are the interaction strength among the nodes. The node size indicates the number of genes represented by the cell type; color chart is proportional to the treatment effects in each cell type. C. Box plot of average cell-cell interaction BiBC (see M&M) for the genes belonging to each treatment effect category (DHA (blue), EPA&DHA (green), EPA (orange) and no category (grey). From the network cell-cell BiBC analysis, genes regulated by DHA, DHA&EPA have higher median BiBC which indicates a higher causal contribution to cell-cell communication than genes regulated by EPA. D. Bar plots for abundance of top BiBC lipids, shown are the PGs (cardiolipin precursors) and SM in NASH preventive study. (Data are mean +/- SD, N=8 mice/treatment group; Ordinary One-way ANOVA, multiple comparisons test with WD+O, *** p<0.005, ****p<0.0001).

**Suppl Figure 3.**
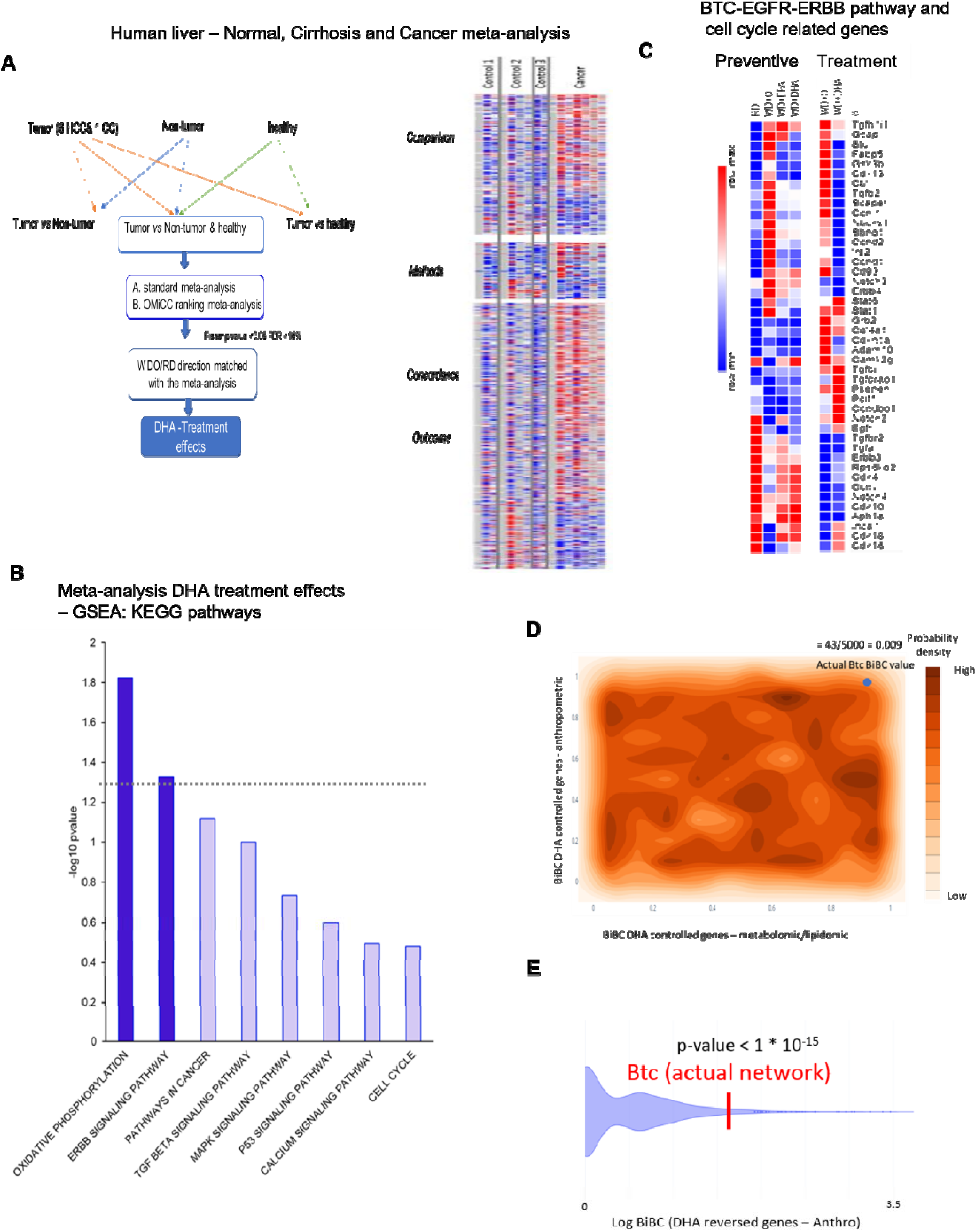
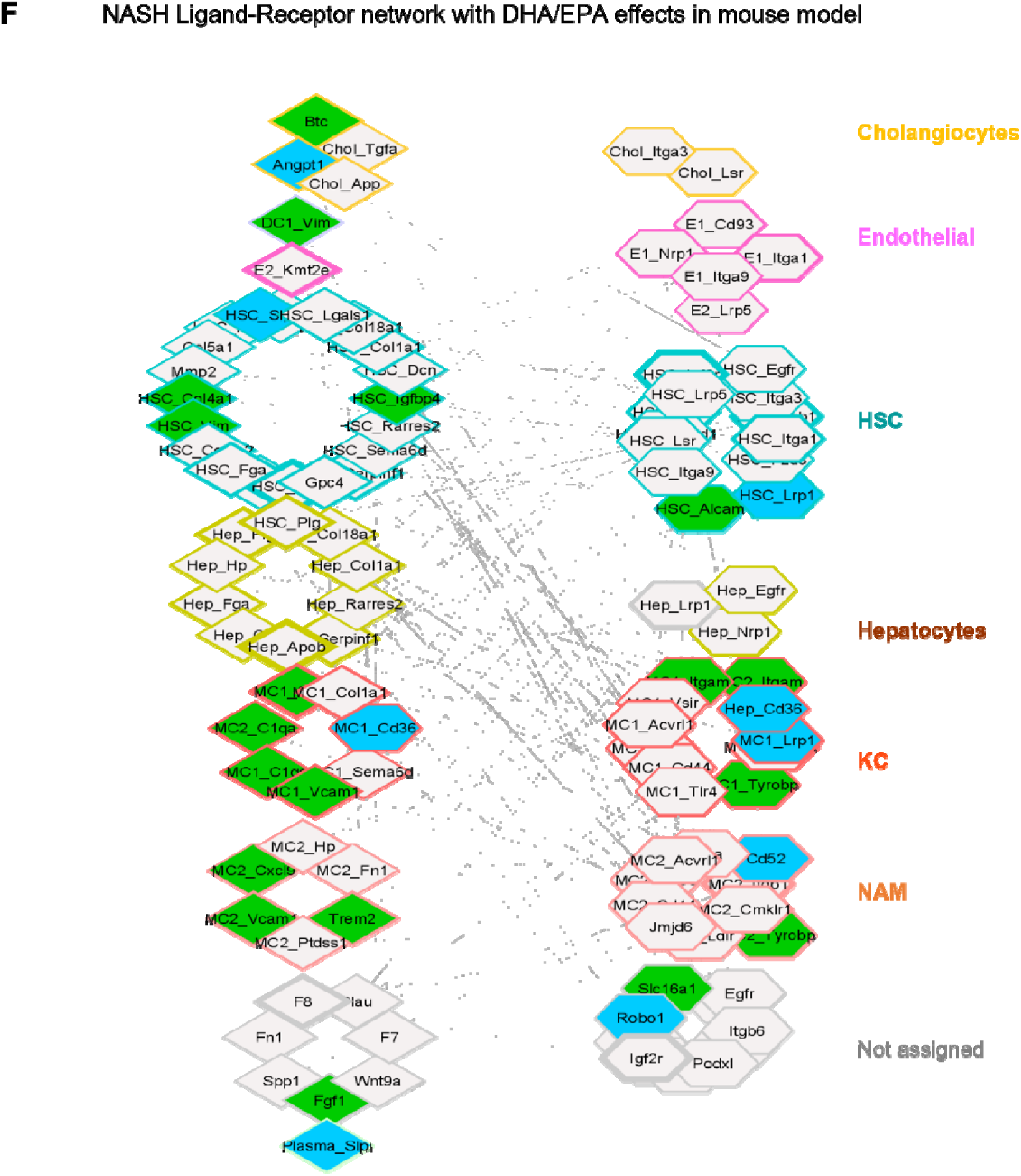
A. Outline for human liver cancer meta-analysis for the orthologous genes representing the NASH mouse model. Right panel shows the heatmap of individual human datasets for selected genes in the meta-analysis (with treatment effects in NASH model). B. The bar graph of gene set enrichment analysis using GSEA (KEGG pathways) for the human cancer meta-analysis genes with DHA effects in mouse models. Data are displayed as −log10(p-value). C. The gene expression for BTC-EGFR-ERBB pathway and cell cycle related genes in the NASH mouse preventive and treatment models. The color scale is indicated from high expression of the genes in red to low in blue. D. Distribution of Btc BiBC values calculated between DHA treatment reversed genes and metabolomic/lipidomic data (x-axis) and DHA controlled genes and anthropometric data (y-axis) in 5000 random networks. Dark regions represent areas where a calculated Btc BiBC value is more likely to be found due to random chance. The probability of finding an actual Btc BiBC value equal to or higher than those seen in the random networks (43/5000) is 0.009. E. Violin plot of the BiBC of Btc between DHA reversed genes and anthropometric data in 5,000 random networks compared to its actual BiBC value in the reconstructed network (one sample Wilcoxon test p < 1 * 10^-15^). F. NASH Ligand-Receptor network with DHA/EPA effects in NASH preventive mouse model. The individual cells expressing the genes inferred from single cell RNA sequence reanalysis are overlayed and cell specific ligand-receptor interactions are shown. Nodes are shown as diamonds (ligands), or hexagon (receptors) and edges are the correlation between the nodes. Node color is according to the treatment effect. The thickness of the node is the cell-cell interaction BiBC, higher the better and is shown thicker. Each cell type and their respective genes are colored differently as labeled. The genes belonging to other cells in addition to specific cells are shown as part of ‘Not assigned’ group.

**Suppl Figure 4.**
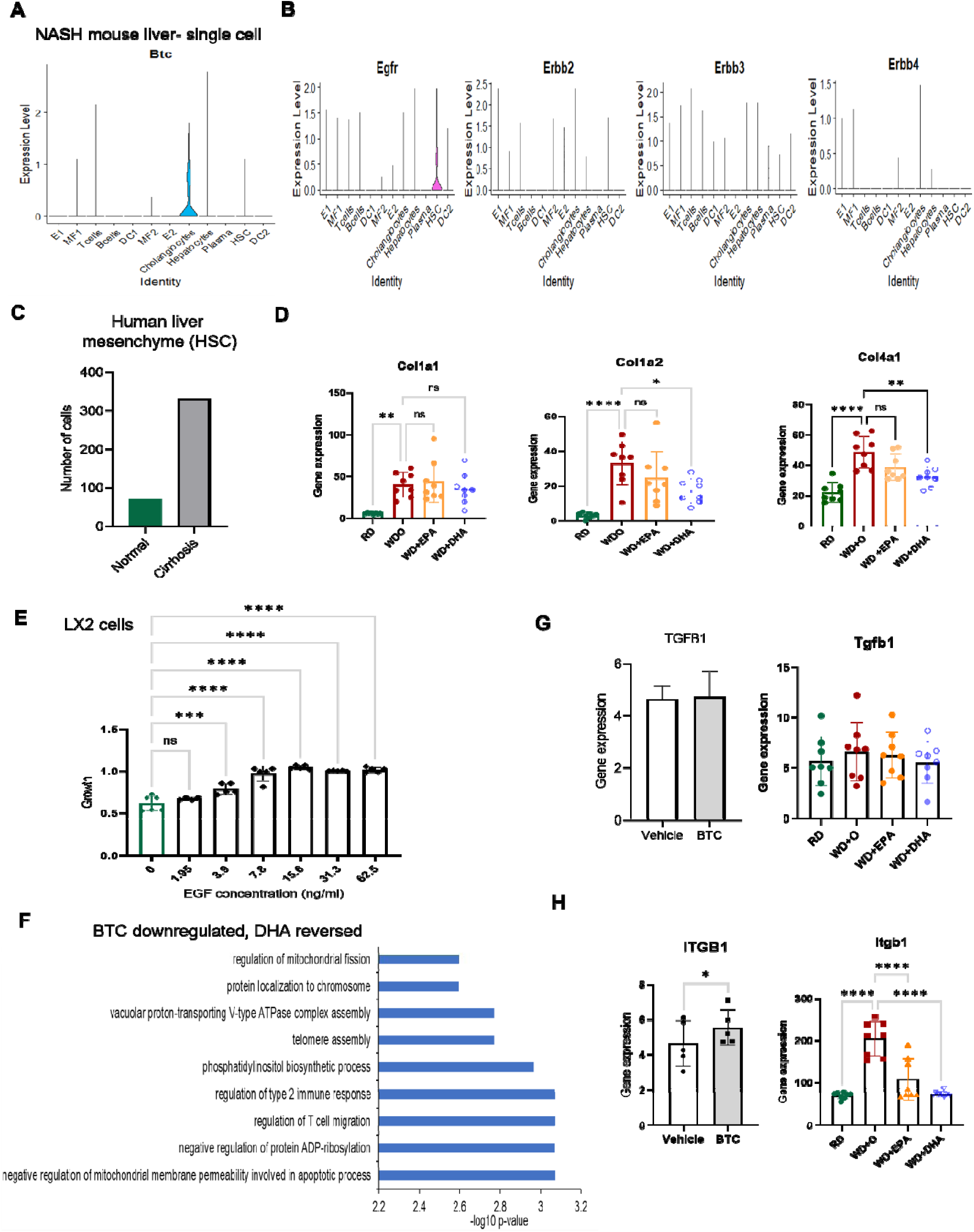
A. The expression of Btc in Mouse NASH liver single cell RNA sequence data shown here with the maximum expression in liver cholangiocytes (MF1-KC; MF2 -NAM). B. The expression of Egfr and other Erbbs in the NASH mouse model single cell RNA sequence data. C. The number of Mesenchymal cells (Hepatic stellate cells) in Human liver samples with enrichment in Cirrhosis with more than 3-fold increase in numbers than normal liver samples. D. The different collagen genes expression in the NASH preventive model shown is in bar graphs colored by treatment effects (Data are mean +/- SD, N=8 mice/treatment group. Ordinary One-way ANOVA, with multiple comparisons test with WD+O, ns (not significant), *p<0.05, ** p<0.001, ****p<0.0001). E. The growth of LX2 cells in response to EGF in a dose dependent manner shown in the bar graph. (Ordinary One-way ANOVA, with multiple comparisons test with Control, ns (not significant), *** p<0.005, ****p<0.0001) F. The gene enrichment analysis shown in a bar plot, regulation of mitochondrial fission and mitochondrial membrane permeability mediated apoptotic pathway are significantly down regulated by BTC treatment in LX2 cells while they are reversed by DHA treatment in the *in vivo* mouse model. G. TGFB1 expression in LX2 cells treated with BTC (grey) (20 ng/ml; N=5 experiments, paired, one-sided t-test, ns (not significant) and in the NASH preventive model is shown in bar graphs colored by treatment effects (Ordinary One-way ANOVA, with multiple comparisons test with WD+O, ns (not significant). H. ITGB1 expression in LX2 cells treated with BTC (grey) (20 ng/ml; N=5 experiments, paired, one-sided t-test, *p<0.05) and in the NASH preventive model is shown in bar graphs colored by treatment effects (Data are mean +/- SD, N=8 mice/treatment group. Ordinary One-way ANOVA, with multiple comparisons test with WD+O, ns (not significant), *p<0.05, ** p<0.001, *** p<0.005, ****p<0.0001).

**Suppl Figure 5.**
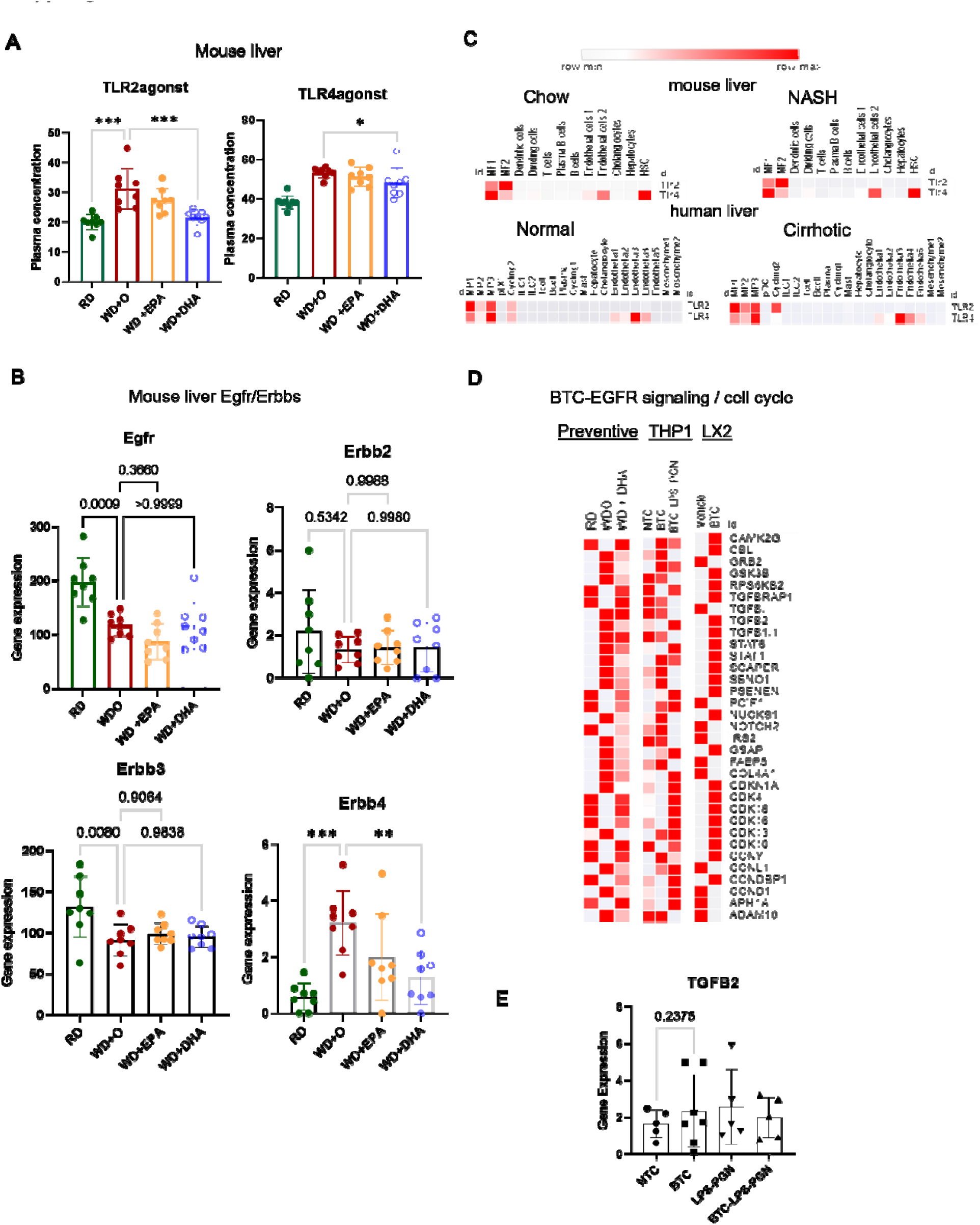
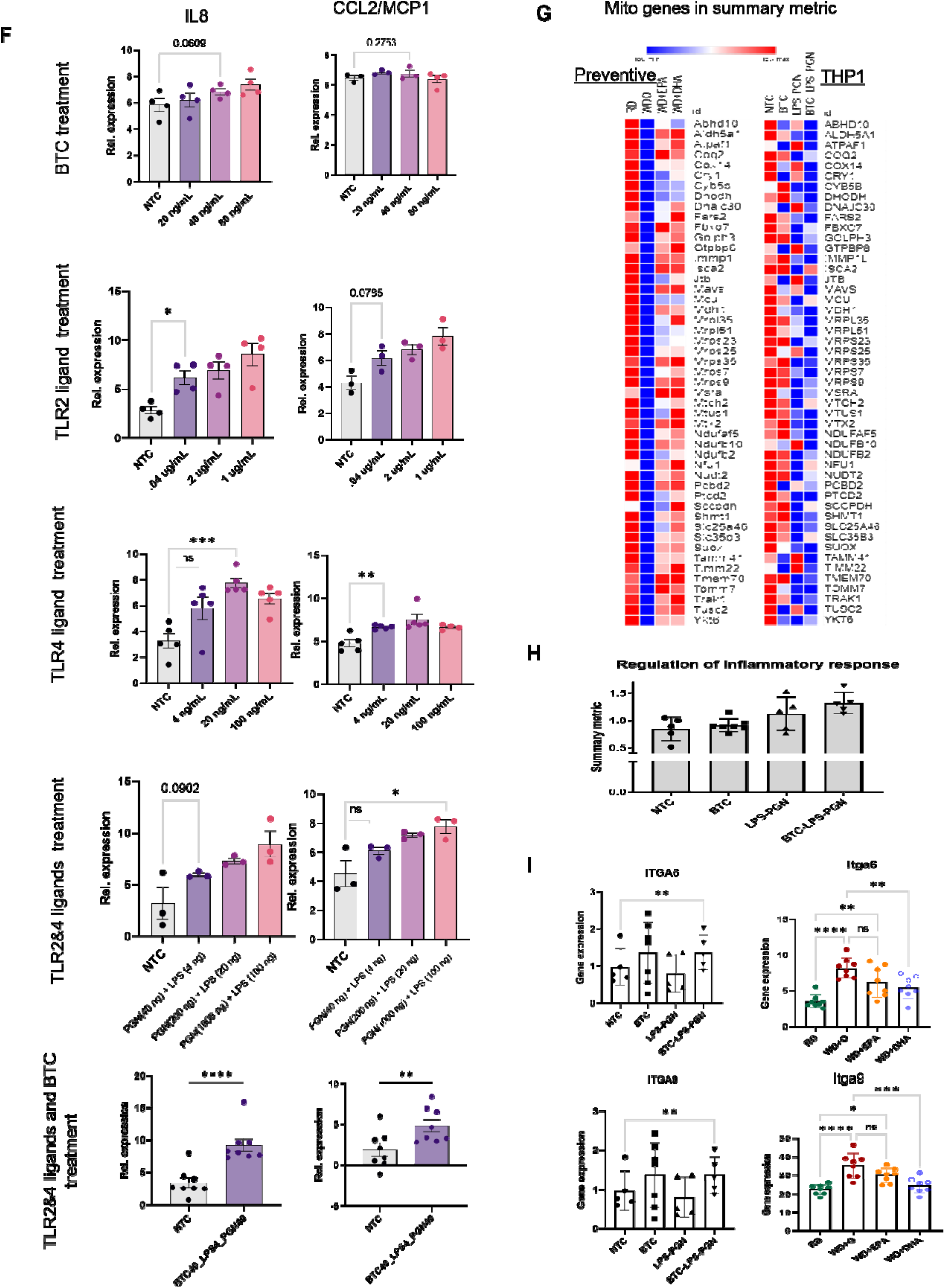
A. Expression of TLR2/4 agonists in the NASH preventive model is shown in bar graphs colored by treatment effects (Data are mean +/- SD, N=8 mice/treatment group). (Ordinary One-way ANOVA, with multiple comparisons test with WD+O, *p<0.05, *** p<0.005) B. The Egfr/Erbbs expression in the NASH preventive model is shown in bar graphs colored by treatment effects (Data are mean +/- SD, N=8 mice/treatment group). (Ordinary One-way ANOVA, with multiple comparisons test with WD+O, ns (not significant), ** p<0.001, *** p<0.005) C. The mouse and human liver (with or without NASH/Cirrhosis) cluster wise average TLR2/4 gene expression from the single cell RNA sequence data. The color scale is indicated from high expression of the genes in red to low in white. D. The gene expression heatmap from BTC/TLR2/4 ligands treated THP-1 and LX2 cells shows genes involved in EGFR pathway and cell cycle pathway that are induced. These set of genes were revered by DHA in the mouse NASH preventive model. The color scale in heatmap is indicated from high expression of the genes in red to low in white. E. Normalized TGFB2 expression in THP-1 cells treated with BTC and or TLR2/4 ligands (5 separate experiments, paired, one-sided t-test, ns (not significant), *p<0.05). F. The dose-response standardization experiments with series of concentrations of BTC, TLR2/4 ligands on THP-1 cells before identifying the lowest concentration for combination of all three together. The well-known cytokines were chosen as markers of gene expression with treatments (IL6 and CCL2; 5-8 experiments, paired, one-sided t-test, ns (not significant), *p<0.05, ** p<0.001, *** p<0.005, ****p<0.0001). G. A heatmap from gene list derived from the enrichment analysis of mitochondria, in summary metric for BTC/TLR2/4 ligand treatment effects in THP1 cells revered by DHA treatment *in vivo* model. The color scale is indicated from high expression of the genes in red to low in blue. H. A summary metric bar graph for BTC/TLR2/4 ligand treatment effects in THP1 cells revered by DHA treatment in vivo model from the enrichment analysis. I. The integrin (ITGA6 and ITGA9) expression in the NASH preventive model is shown in bar graphs colored by treatment effects (Data are mean +/- SD, N=8 mice/treatment group). (Ordinary One-way ANOVA, with multiple comparisons test with WD+O, ns (not significant), *p<0.05, ** p<0.001, *** p<0.005, ****p<0.0001) and in THP-1 cells treated with BTC and or TLR2/4 ligands (N=5 experiments, paired, one-sided t-test, ns (not significant), *p<0.05).

## Materials and Methods

### Animals and diets

Study design for DHA mediated NASH prevention and remission in Male *Ldlr ^-/-^* mice. This study was carried out in strict accordance with the recommendations in the Guide for Care and Use of Laboratory animals of the National Institutes of Health. All procedures for the use and care of animals for laboratory research were approved by the Institutional Animal Care and Use Committee at Oregon state University (Permit number: A3229-01). Anthropometric, plasma and liver samples used in this study were obtained from our previously published NASH prevention and treatment studies (Depner *et al*., 2013a; Lytle *et al*., 2015). Briefly, male mice (B6:129S7-*Ldlr^tmHer^*^/j^, stock# 002207) were purchased from Jackson Labs and were group housed (4 mice/cage; n=8 mice per group) and maintained on a 12-hour light/dark cycle. Mice were acclimatized to the Oregon State University Linus Pauling science center vivarium for 2 weeks before proceeding with the experiments.

#### NASH Prevention Study

This study was designed to determine if EPA and DHA differed in their capacity to prevent western diet-induced NASH (Depner *et al*., 2013a). Mice consumed the one of the following 5 diets, ad libitum for 16 weeks; each group consisted of 8 male mice. Purina chow 5001 consisting of 13.5% energy as fat and 58.0% energy as carbohydrates was used as the Reference diet (**RD**). The western diet (**WD**) was obtained from Research Diets (12709B) and used to induce NASH. The WD consisted of 17% energy as protein, 43% energy as carbohydrate, and 41% energy as fat; cholesterol was at 0.2% wt:wt. The WD was supplemented with olive oil (**WDO**), eicosapentaenoic acid (**WD + EPA**), docosahexaenoic acid (**WD + DHA**), or both EPA and EHA (**WD + EPA + DHA**). EPA and DHA were added to the diets to yield 2% of total calories; for the EPA + DHA, each was added to yield 1% of total calories, i.e. 2% total calories as C_20-22_ ω3 PUFA. Olive oil was added to the WD to have a uniform level of fat energy in all the WDs. Preliminary studies established that the addition of Olive oil had no effect on diet-induced fatty liver disease in *Ldlr^−/−^* mice. EPA was purchased from Futurebiotics as Newharvest EPA, a DuPont product, while DHA was obtained as a generous gift from DSM, formally Martek Bioscience). The amount of EPA or DHA added to the diets is equivalent to the amount prescribed for treating patients for hypertriglyceridemia (Davidson *et al*, 2007; Harris *et al*, 1997). At the end of the 16-week feeding trial, mice were euthanized with CO2, blood (RBC and plasma) and liver were collected and stored at −80°C for later extractions of RNA, lipids, proteins.

#### NASH Treatment Study

This study was designed to assess the capacity of DHA to reverse the effects of WD-induced NASH (Lytle *et al*., 2017). As such, male *Ldlr* ^-/-^ mice were fed the WD for 22 wks. These mice were separated into 2 groups: one group was maintained on a WD + olive oil, with the other group was maintained on the WD + DHA. The diet composition was as described above for the prevention study. Both groups were euthanized after 8 wks of these diets. A control group was maintained on the RD for 30 wks. At the end of the study, mice were euthanized, and samples collected and processed as above.

### Cell culture

***LX2*** cells were obtained from SL Friedman (Mount Sinai Medical School)(Xu *et al*., 2005). LX2 cells are activated human hepatic stellate cells; they were maintained in DMEM with 10% FCS containing penicillin and streptomycin. Cells were plated onto 100 mm plastic petri dishes ∼100,000 cells/plate and treated with fatty acids (at 50 μM) in endotoxin-free BSA (at 20 μM) during the growth phase. Fatty acids [oleic acid (18:1 ω9) and DHA (22:6, ω3); Nu-Chek Prep] were used at 50 μM for 2- 48-hour cycle treatments. After this pre-treatment, cells were trypsinized, washed in PBS, counted and plated at 3000 cells/well in a 96-well plate. Cells were fed DMEM with 1 % FCS without or with betacellulin (human recombinant betacellulin (BTC), R&D Systems) for 72 hrs; the concentration ranged from 1.25 to 25 ng/ml. At the completion of BTC treatment, media was removed, washed with PBS and DNA/well was quantified using the CyQuant cell proliferation assay (Thermofisher); DNA fluoresces was quantified [excitation at 485 nm & emission at 530 nm. This experiment was repeated 3 times.

#### Collagen Production

Pico Sirius red quantitation of collagen production: Cells were pretreated with fatty acids as described above, trypsinized, counted and plated onto 12-well cell culture plates at 80,000 cells/well in DMEM +1% FCS and treated without and with 20 ug/ml BTC for 72 hrs. At the end of treatment, media was removed, cells were washed with PBS and stained with pico Sirius Red (Abcam) for 2 hrs at room temperature. After staining cells were washed with an acetic acid solution, photographed and the stain was removed using 50 mM NaOH. The level of staining per well was quantified at 540 nm. The level of protein/well was quantified after solubilizing the protein in 50 mM NaOH and using the Pierce BSA kit. This experiment was repeated 4 times.

#### Cell Vitality

Alamar Blue (Creative Labs) assessment of cell vitality (NADH conversion to NAD^+^): Using the protocol describe above, we assessed the vitality of cells after treatment with fatty acids and BTC. Alamar blue (50 μl/1.0 ml media) was added to the cells and fluorescence was measured at 590 nm 4 hrs after Alamar Blue addition.

#### RNASeq

RNA was extracted from LX2 cells using Trizol (Invitrogen) as previously described (Lytle *et al*., 2015) from cells treated with fatty acids (oleic acid and DHA at 50 μM for 72 hrs) as described above. Cells were seeded onto 6-well plates at 40,000 cells/well; and cells were treated with fatty acids for 96 hrs as describe above. Afterward, media was removed, and cells were treated without and with BTC at 20 ng/ml for 72 hrs. Cells were harvested for RNA extraction (Triazol). cDNA was prepared for RNASeq analysis as described. *THP-1 cells*

### BTC and TLR co-stimulation

THP-1 monocytes were cultured in RPMI 1640 medium adjusted to contain 4.5 g/L glucose and supplemented with 1% Penicillin/Streptomycin, 10% FBS, 1 mM sodium pyruvate, 10 mM HEPES. For experiments, monocytes were first seeded in 24 well plates at 400,000 cells/well in 1 mL of complete medium with 50 ng/mL PMA (phorbol 12-myristate 13-acetate) to induce polarization, for 24 hours. Then, attached cells were washed with sterile PBS to remove residual serum and PMA containing media. Cells were stimulated with a TLR2 agonist at 40 ng/mL (PGN-BS, Invivogen, San Diego, CA), TLR4 agonist at 4 ng/mL (LPS-B5, Invivogen, San Diego, CA), BTC at 40 ng/mL (human recombinant Betacellulin protein, R&D systems, MN), or combinations of all for 6 hours. Treatments were prepared in serum-free semi-complete RPMI 1640 media (see above for other components) supplemented with 20 μM BSA and 50 μM BHT (Butylated hydroxytoluene). Cells were lysed with 300 μL RLT buffer (Qiagen) and cell lysates were stored at −80 °C.

### Primers

#### qRT-PCR data analysis

THP-1 cells’ response to TLR and BTC stimulation was assessed by qRT-PCR. Briefly, raw Cycle Threshold (CT) values from the StepOnePlus Real Time PCR instrument for genes of interest were normalized to CT values of a housekeeping gene, TMEM59, by delta CT method and relativized by 2_−_ΔCT. Data were then median normalized and Log_2_ transformed before being plotted in GraphPad Prism 9.

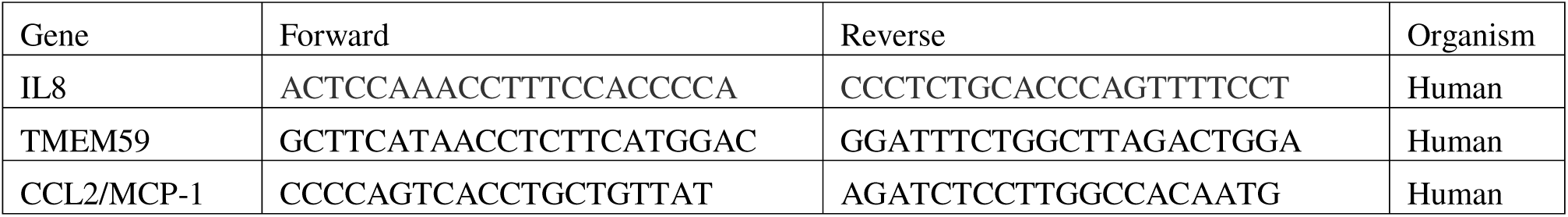

### RNA extraction and RNA sequencing library preparation

RNA was extracted from cell lysates using the RNeasy Mini Kit and subjected to a DNase treatment according to manufacturer’s protocols (Qiagen) then stored at −80 °C until further use. mRNA libraries were prepared for sequencing with the Lexogen QuantSeq 3’ mRNA-Seq Library Prep Kit (FWD) HT for Illumina Sequencing platforms (Kit#k15.384) and sequenced on the Illumina NextSeq at Oregon State University.

### cDNA and qRT-PCR

cDNA was prepared from 0.5-1ug of RNA via reverse transcription using the qScript XLT cDNA SuperMix kit (Quantabio). qRT-PCR was performed for gene expression using the AzuraView GreenFast qPCR Blue Mix HR (Azura Genomics). 96-well plates were prepared with 10 ng of cDNA in triplicate reactions for each gene and sample and run on an Applied Biosystems StepOnePlus Real Time PCR instrument.

### Sequencing of RNA (RNAseq)

RNA libraries were prepared with the QuantSeq 3’mRNA-Seq Library Prep Kit (Lexogen) for the Apollo 324 NGS Library Prep System and sequenced using Illumina NextSeq. The sequences were processed to remove the adapter, polyA and low-quality bases by BBTools (https://jgi.doe.gov/data-and-tools/bbtools/) using bbduk parameters of k=13, ktrim=r, forcetrimleft=12, useshortkmers=t, mink=5, qtrim=r, trimq=15, minlength=20. Then the reads were aligned to mouse genome and transcriptome (ENSEMBL NCBIM37) using Tophat (v2.1.1) with default parameters. The number of reads per million for mouse liver genes were counted using HTSeq (v 0.6.0) and quantile normalized. Cell lines sequencing/analysis was performed similarly, but with the bbduk parameters of k=13, ktrim=r, forcetrimleft=11, useshortkmers=t, mink=5, qtrim=r, trimq=10, minlength=20. The reads were aligned to the human genome and transcriptome (Gencode v40) using STAR v2.5.3a. BRB-ArrayTools was used to identify differentially expressed genes between treatments.

### Metabolomes and Lipidomes

Data were prepared, and analysis were carried out from the NASH Prevention study and Treatment study. Hepatic lipids were extracted and subject to UPLC/MS/MS analysis as previously described (Garcia-Jaramillo *et al*, 2019) with minor modifications.

### Treatment categorization of omics data

The genes and other parameters whose expressions or values are significantly changed by WD (FDR<10%) were considered for treatment effects. Out of these parameters and genes, those that have treatment effect reversal with a P-value<0.05 were categorized as DHA if uniquely DHA (opposite to WD+O/ND & WD+DHA/WD+O P-value<0.05), uniquely EPA (opposite to WD+O/ND & WD+EPA /WD+O P-value<0.05), similar in both EPA&DHA (opposite to WD+O/ND & both WD+EPA /WD+O and WD+DHA/WD+O, P-value<0.05) and with no treatment effect (NA; with both WD+EPA /WD+O and WD+DHA/WD+O, P-value>0.05). To be consistent with many previous studies, the treatment effects were also tested in the combination of preventive treatment though not used further in the analysis (EPA+DHA; WD+EPA+DHA /WD+O, P-value<0.05).

### Reconstructing the NASH liver multi-omic network

The network reconstruction was carried out as described in the previous papers from our group with minor dataset specific modifications (Dong *et al*., 2015; Li *et al*., 2022). The genes and other parameters whose expressions or values are significantly changed by WD (FDR<10%) were chosen for constructing the NASH network. First, from liver tissue between all pairs of genes (GE) and metabolic parameters (phenotypes, P) spearman rank correlations were calculated by pooling the samples per diet (WD+O, WD+EPA and WD+DHA). Meta-analysis was performed to retain edges with same sign of correlation coefficient in all three diets. Edges were furthered filtered by the following criteria: individual p-value of correlation within each diet from pooled experiments <20%, combined Fisher’s p-value over diets from pooled experiments <5% and FDR cutoff of 10% for edges within tissues and for phenotypes and between lipidomic, metabolomic and plasma biochemicals and edges needed to satisfy principles of causality [i.e., satisfied fold change relationship between the two partners in the WD+O vs. ND comparison]. Next, correlations were calculated per diet for the experiment pairwise between parameters (Gene Expression + Lipidomic and Metabolomic data (LM)) and (P+LM). Finally, edges obtained from pooling were retained if they had the same sign of correlation coefficient as in 3 groups (3 diets, WD+O, WD+EPA and WD+DHA). False positive edges while pooling the different diets were removed in the creation of the network. The proportion of genes, metabolites and lipids that made it to the final network (following statistical cutoffs) was determined after applying selection for significance in differentially expressed parameters in liver prior to applying correlation cutoffs.

### Single cell RNA sequence data analysis

#### Datasets

Single cell dataset (GSE129516) for mouse NASH model was obtained from single cell RNA-sequence on non-parenchymal cells of healthy vs NASH mouse liver. These are then reanalyzed and used in our multi-omic network analysis to infer liver cell types. Human liver single cell RNA sequencing data (GSE136103) from normal and Cirrhosis patients were reanalyzed (Ramachandran *et al*., 2019; Xiong *et al*., 2019).

#### scRNA sequence analysis

The raw gene expression matrix (UMI counts per gene per cell in the liver tissue) was filtered, normalized, and clustered using a standard Seurat version 3.1.0 in R [https://www.R-project.org/] (Stuart *et al*, 2019). Cell and gene filtering were performed as mentioned in previous publications. During quality filtering, cells with a very small library size (<5000) and a very high (>12%) mitochondrial genome transcripts were removed. The genes detected (UMI count > 0) in less than three cells were also removed from further processes. Then log normalization and further clustering is performed using standard Seurat package procedures. Principal component analysis was used to reduce dimensions that represent cell clusters. The number of components from this analysis used for the elbow of a scree plot, which further aid in selecting the significant clusters. The different cell type clusters in a sample were visualized using t-distributed Stochastic

Neighbor Embedding of the principal components as implemented in Seurat. The liver tissue specific cell-type identities for each cluster were determined manually using a compiled panel of available immune and other cell specific marker expression as per the previously published papers (Ramachandran *et al*., 2019; Xiong *et al*., 2019).

### Single cell RNA sequence for assignment of gene in the NASH network to a specific cell type

The normalized UMI > 1.0, with a fold change significantly and uniquely expressed genes in a cell specific cluster were then assigned to that specific set of genes in the network to indicate they belong to that specific cell type. It is the primary rule to assign a gene to a cell type. Next, higher expression in the cell cluster (and optional fold change (log2FC >0.25, p value <0.05) for a gene gets assigned with that specific cell type of the tissue. Here, additionally, ranking by average expression for each gene in the clusters helps to determine its cluster specificity by the higher expression in that cell type than another in the whole tissue. This is implemented for evaluating highest cell cluster average expression of a gene, among all other cell clusters in network.

### Detecting subnetworks and functional enrichment

Infomap (mapequation.org) was used to identify subnetworks using the default commands. Functional enrichment of clusters was then performed using metascape (http://metascape.org) (Zhou *et al*, 2019).

Additionally single cell data overlay on NASH network as mentioned above allowed the cell type specific gene sub clusters.

### Identifying of key nodes between subnetworks using BiBC analysis

Analysis of networks was performed using the python package NetworkX v2.2. Bipartite betweenness centrality (BiBC) was calculated between all cell cluster pairs (66 in total) of the twelve clusters previously identified based on single cell data overlay on NASH network. The nodes were then ranked by their resulting BiBC and scaled to range of 0-100. BiBC was also calculated and scaled similarly pairwise between all genes and anthropometric data, between all genes and Lipidomic/Metabolomic data.

### Creation and analysis of random networks

Random networks were created similar to as was described in Kahalehili et al., 2021 (Kahalehili *et al*, 2020). Briefly, 5,000 Erdos-Renyi random networks were created, utilizing the same number of nodes and edges that were present in the real, reconstructed network. BiBC was calculated both 1) between DHA controlled genes and anthropometric nodes and 2) between DHA controlled genes and DHA controlled metabolomic/lipidomic data. Plotly (https://chart-studio.plotly.com/create/#/) was used to create the 2D contour histograms for visualization of the random network results.

### Intrahepatic ligand-receptor interaction network

The knowledgebase of ligand receptor interaction information available for mouse genes (Abugessaisa *et al*., 2021), was overlayed on to reconstructed NASH Multiomics network genes. Additionally, this allowed the interrogation of ligand-receptor network with respect to each cell in the network that a gene can be represented as ligand or receptor.

### Human liver cancer meta-analysis

Human cancer data from hepatic cellular cancer (HCC) and cholangiocytes cancer (CC) were selected from GEO data sets (GSE14520, GSE26566, GSE102079, GSE56140, GSE98617, GSE76427, GSE84005). Meta-analysis using RankProd method described in the publicly available tool OMiCC (Shah *et al*, 2016) was carried out for these 7 sets of data with 32 healthy, 260 non-tumor samples and 544 tumor samples (Suppl Figure 3A). At first sample sets of both tumor types were compared against their respective paired non-tumor or healthy samples.

Additionally, a standard (Fisher) approach of meta-analysis was also carried out and an overlapping set of genes were selected.

Then a signature set of genes were identified by matching (among orthologous) genes between mouse and human using Biomart (Cunningham *et al*, 2022) for the concordant fold change direction as WD+O/RD in the NASH network model and genes that are significant with FDR less than 15% and Fisher p value less than 5%. The human liver cancer meta-analysis signature genes were overlapped with the NASH mouse model network genes with treatment effects to identify subset of genes.

### Gene Ontology analyses

The gene ontology and functional enrichment were carried out using Metascape and innateDB (Breuer *et al*, 2013; Zhou *et al*., 2019) with mouse or human reference databases. The reference database is determined depending on the species of the specific data in question.

### Pathway summary metric

Using published approach (Levine *et al*, 2006) with minor adaptions described in our previous papers (Koscso *et al*, 2020; Shulzhenko *et al*, 2011), each top gene ontology pathway enriched in the THP-1 experiment (with BTC and TLR ligands and significantly revered by DHA in NASH preventive model) was recognized and the genes belonging to the pathways were identified.

Using median normalization, a value for each gene in each treatment replicate and then additionally using their median, summary metric was calculated for the individual pathway. A paired t-test between normal and treatment condition were performed to evaluate significance.

### Statistical analysis

In mouse studies data are expressed as geometric mean of replicates. Data are shown as the mean ± standard deviation in animal studies and *in vitro* experiments. Group comparisons were performed using an unpaired *t* test and Ordinary One-way analysis of variance (ANOVA), followed by Tukey’s *post hoc* or Dunnett’s multiple comparisons tests, where *p* <0.05 indicates statistical significance. In cell line and RNA sequence analysis, comparisons between groups were performed using Student’s *t* test or the Mann-Whitney *U* test and Kruskal-Wallis test when appropriate. Categorical variables are shown as counts and percentages. Differences between categorical variables were assessed with the chi-squared test or Fisher’s exact test. Spearman’s rank correlation rho coefficients were calculated for network edges between all parameters using R statistical packages. Details of statistical analyses are described additionally in the corresponding figure legends. GraphPad Prism 9 was used for all analyses.

